# Single-AAV-mediated cardiomyocyte-specific adenine base editing ameliorates cardiac hypertrophy in vivo

**DOI:** 10.64898/2026.11.12.765297

**Authors:** Thomas Monnikhof, Su Ji Han, Laura Stumpo, Serj Koshian, Ceren Agirman, Hesther de Ruiter, Alessandra Luppino, Tessa Wigman, Margot Chevalier, Ilaria Perini, Danielle Versteeg, Martijn Wehrens, Anne E. de Leeuw, Maya Wright-Clark, Sebastiaan J. van Kampen, Lorena Zentilin, Danny D. Sahtoe, Prasad R. Konkalmatt, Mohammad A. Mandegar, Christopher A. Reid, Kathryn N. Ivey, Mauro Giacca, Cornelis J. Boogerd, Eva van Rooij

## Abstract

Hypertrophic cardiomyopathy (HCM) is the most common inherited cardiac disorder, frequently accompanied by cardiac hypercontractility, diastolic dysfunction, heart failure and sudden cardiac death. The disease is predominantly caused by pathogenic variants in genes encoding sarcomeric proteins and current therapeutic strategies largely focus on symptom management. Adeno-associated virus-based (AAV) delivery of base editing tools has emerged as an alternative strategy that acts directly on the genetic cause of disease. However, canonical SpCas9-based editors are too large to package into a single-AAV vector which mandates dual-AAV approaches that require higher overall vector doses with elevated toxicity risk and increased manufacturing costs. Here, we report the development and application of a single vector approach for cardiomyocyte-specific adenine base editing to correct a Dutch founder mutation in *MYBPC3* (c.2827C>T, p.Arg943Ter) that causes HCM. We systematically compared compact base editors in human induced pluripotent stem cell-derived cardiomyocytes (hiPSC-CMs) and selected the most efficient configuration, which reversed disease-associated hypercontractility in engineered human myocardium (EHM). To fit the editor/guide construct in a single-AAV vector, we screened a library of rationally designed small chimeric promoters and identified 6C2 to effectively drive expression in hiPSC-CMs and cardiac tissue of both mice and non-human primates (NHP) with exceptional specificity. Delivered in a single next-generation cardiotropic AAV capsid, we achieved therapeutic correction in an HCM mouse model at doses 20-fold below published dual-AAV cardiomyopathy base editing approaches. These results establish cardiomyocyte-specific, single vector-mediated base editing as a viable therapeutic strategy for inherited cardiomyopathies.

## Introduction

Hypertrophic cardiomyopathy (HCM) is the most common inherited cardiac disorder, affecting approximately 1 in 200-500 individuals worldwide^1^. It is characterized by unexplained left ventricular hypertrophy, frequently accompanied by diastolic dysfunction, heart failure, an increased risk of sudden cardiac death (SCD), and is typically inherited in an autosomal dominant manner with variable penetrance and phenotypic expression^2,3^. HCM is commonly associated with hypercontractility, which can contribute to dynamic outflow tract obstruction and impaired ventricular function. The disease is predominantly caused by pathogenic variants in genes encoding sarcomeric proteins, the fundamental contractile units of cardiomyocytes^4^. Current therapeutic strategies for HCM largely focus on symptom management and prevention of SCD rather than on the underlying genetic cause. The recent approval of the myosin inhibitor mavacamten marked an important advance for obstructive HCM, providing clinical benefit through selective, reversible inhibition of β-cardiac myosin that reduces sarcomeric hypercontractility and improves ventricular compliance^5^. However, in the recently completed phase 3 ODYSSEY-HCM trial, mavacamten did not significantly improve peak oxygen uptake or symptom burden in patients with non-obstructive HCM, underscoring that pharmacological modulation of contractility does not correct the underlying genetic lesion and leaves a substantial fraction of patients without an effective disease-modifying therapy^6^.

AAV-based gene therapy and genome editing have emerged over the past decade as complementary strategies that act directly on the genetic cause of disease^7,8^. Gene replacement therapy for MYBPC3 has demonstrated clear efficacy in *Mybpc3*-deficient mouse models^9,10^. More recently, an early-phase, first-in-human trial of an AAV9-delivered *MYBPC3* transgene yielded encouraging interim safety and efficacy signals, including reductions in left ventricular mass and improvements in clinical symptom scores^11^. However, gene replacement therapy relies on life-long expression from an episomal AAV genome in post-mitotic cardiomyocytes. Whether this expression can be therapeutically adequate over a patient’s lifetime remains unresolved^12^. Additionally, conventional gene supplementation is not suitable for patients harboring dominant-negative mutations or toxic, truncated peptides, which require direct correction at the genomic level. Genome editing offers a fundamentally different, potentially one-time curative strategy: rather than supplementing wildtype protein, it permanently corrects the pathogenic variant itself^13^. Among current genome editing technologies, base editing is particularly attractive because it can install single-nucleotide substitutions without generating double-strand DNA breaks, by tethering a Cas9 nickase to a nucleotide deaminase that catalyzes base conversion within a defined editing window^14,15^.

Base editing has already shown therapeutic promise in mouse models of sarcomeric HCM. Dual-AAV9 delivery of a split-intein adenine base editor durably corrected a *MYH7* HCM variant in vivo^16,17^. In a separate study, AAV9-delivered SpRY-Cas9-based ABE8e correction of a *Mybpc3* nonsense allele in mice was shown to outperform AAV-mediated *Mybpc3* gene replacement therapy^18^. These studies demonstrate that base editing can durably correct sarcomeric HCM variants in vivo.

The requirement for a dual-AAV strategy arises because current SpCas9- or SpRY-Cas9-based base editor sequences exceed the practical packaging capacity of a single-AAV vector (∼4.7 kb)^19^. This is solved by dividing editor fragments over two vectors which are reconstituted through trans-splicing inteins once both have entered the same cell^20,21^. To achieve therapeutic efficacy, dual-AAV strategies mandate significantly higher total viral vector doses, with a corresponding increase in toxicity risk, creating an important translational limitation. High systemic AAV doses have been associated with severe and, in some cases, fatal adverse events, including complement activation, thrombotic microangiopathy, and acute multiorgan toxicity^22–24^. A single-AAV strategy would therefore simplify design and manufacturing, and could potentially achieve therapeutic editing at substantially lower total viral doses, hence offering a simpler regulatory and clinical development path with an improved safety margin.

Beyond size, canonical base editors are further constrained by strict protospacer-adjacent motif (PAM) requirements that limit the range of targetable pathogenic variants. Near-PAMless Cas9 variants, including SpRY, address this limitation by recognizing a broader range of PAM sequences, including many NRN and NYN motifs^25^. However, increased PAM flexibility also expands the number of potential binding sites throughout the genome relative to wildtype SpCas9. This includes increased genome-wide off-target editing, more frequent chromosomal translocations, and markedly higher rates of double-stranded breaks (DSBs)^26–28^. Hence, expanding targetability through PAM relaxation may introduce safety concerns that are particularly important for therapeutic in vivo applications.

In this study, we set out to develop a cardiomyocyte-specific base editing strategy compatible with single-AAV vector delivery to correct the Dutch founder *MYBPC3* nonsense mutation (c.2827C>T, p.Arg943Ter) that causes HCM^29^. Although most patients are heterozygous, homozygous or compound heterozygous carriers develop fatal, early-onset cardiomyopathies within the first months of life^30^. Crucially, this target locus lacks a suitably positioned canonical NGG-PAM to align the pathogenic mutation within the catalytic window of conventional SpCas9-derived base editors, necessitating alternative Cas9 nickases with distinct PAM recognition domains. We systematically compared compact Cas9 variants, deaminases and single guide RNA (sgRNA) designs in both human and mouse models of HCM to identify the most efficient configuration. Driven by the stringent packaging capacity of AAVs, we additionally screened a library of rationally designed short (250 bp) chimeric promoter candidates in hiPSC-CMs in vitro and evaluated the top-performing candidates in non-human primates (NHPs) and mice in vivo. Next, we compared 6C2, our most highly active candidate in NHP and mouse hearts, to other short promoters and confirmed 6C2 as an efficient cardiomyocyte restricted promoter for single-AAV-mediated base editor expression. Finally, by packaging all editor components in a single next-generation cardiotropic AAV capsid, we achieved therapeutic benefit in an HCM mouse model using doses 20-fold lower than those in previous cardiac base editing studies. Together, our findings establish a single vector compatible base editing strategy for a clinically relevant HCM variant and demonstrate the therapeutic potential of compact, tissue-restricted base editing for the treatment of inherited cardiomyopathies.

## Results

### Base editing corrects the human *MYBPC3* R943X founder variant using a compact Cas9 variant

To evaluate adenine base editing (ABE) as a strategy to correct HCM-causing *MYBPC3* variants in cardiomyocytes, we used hiPSCs harboring the *MYBPC3* c.2827C>T (R943X) mutation (**Fig. 1a**). Previous studies have shown that patient-derived hiPSC-CMs heterozygous for this mutation maintain their contractile function and MYBPC3 protein levels comparable to isogenic controls^31^. Therefore, we focused on homozygous *MYBPC3^R^*^943^*^X/R^*^943^*^X^* hiPSC-CMs to establish a system in which the molecular and functional consequences of base editing on MYBPC3 loss could be assessed. Homozygous *MYBPC3^R^*^943^*^X/R^*^943^*^X^* hiPSC-CMs showed markedly reduced *MYBPC3* transcript levels compared to *MYBPC3^WT/Corr^* hiPSC-CMs, while full-length MYBPC3 or the predicted truncated MYBPC3 protein remained undetectable (hereafter: R943X and Corr, respectively; **Fig. 1b-e**).

**Fig. 1:**
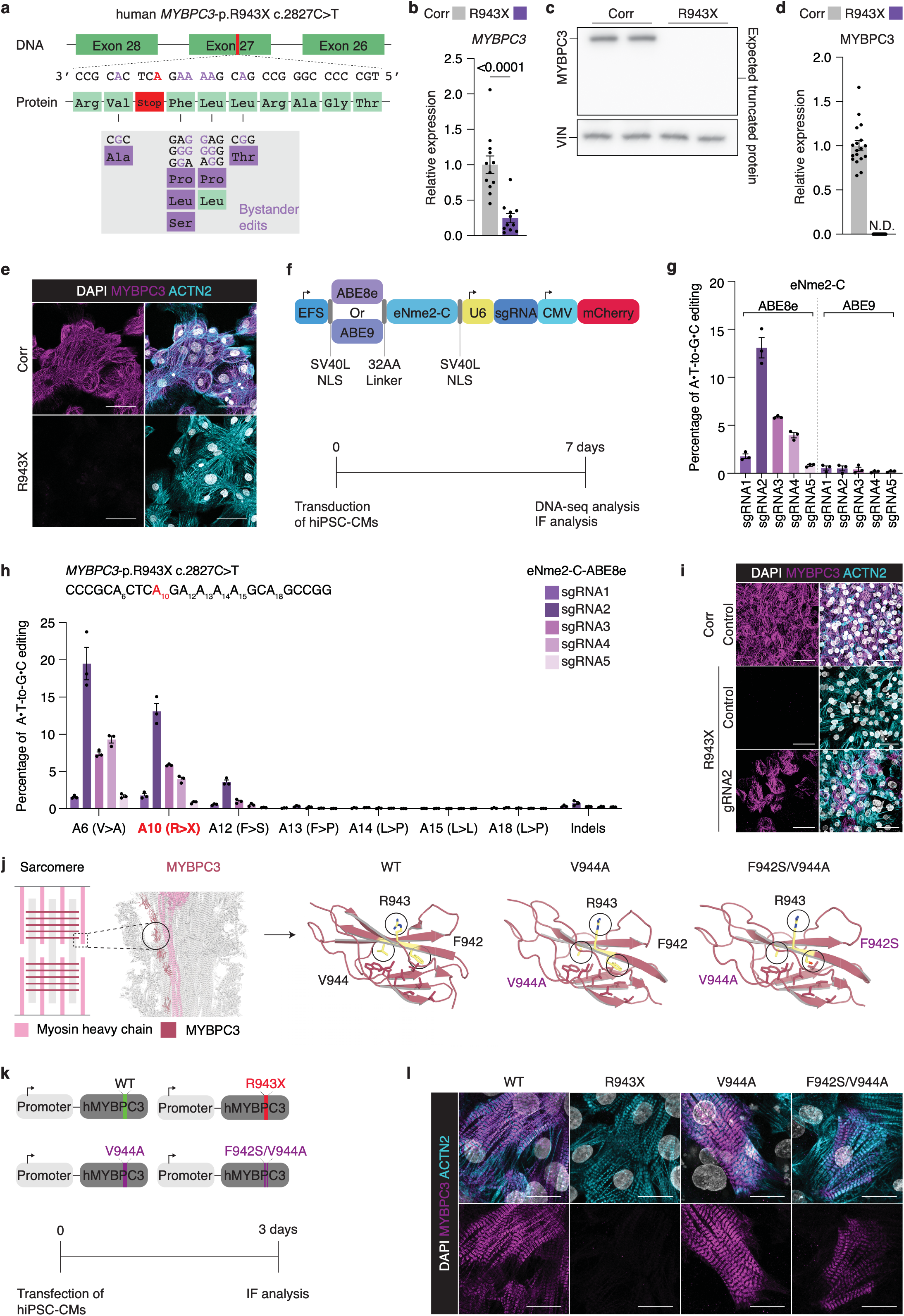
Base editing corrects the pathogenic HCM-causing *MYBPC3*-R943X c.2827C>T variant and restores protein in hiPSC-CMs. **a,** Schematic of the R943X-containing genomic region (exon 27, red). The sequence is shown 3′ to 5′ to indicate target adenines within the protospacer. Potential bystander edits are shown in purple with corresponding amino acid changes. **b,** qPCR analysis of *MYBPC3* expression in isogenic control (Corr) and R943X hiPSC-CMs. Expression was normalized to *GUS* and *HARP* using the ΔΔCt method. Primers are listed in **Table S1**. Data are from two independent cardiomyocyte differentiations with 5-6 replicates. **c,** Representative MYBPC3 Western Blot with quantification in **d**. **d,** Quantification of MYBPC3 expression normalized to VIN across three independent cardiomyocyte differentiations (*n*=6 technical replicates). **e,** Representative MYBPC3 immunofluorescence images of Corr (top row) and R943X hiPSC-CMs (bottom row). Scale bar: 50 *μ*m. DAPI, white; MYBPC3, magenta; ACTN2, cyan. **f,** Schematic of all-in-one lentiviral editing constructs. ABE8e or ABE9 was fused to eNme2-C and driven by the EFS promoter, followed by a U6-sgRNA cassette. A CMV-mCherry cassette enabled visualization of transduction. **g,** Editing efficiency by HTS data at the target adenine (dots represent *n*=3 independent cardiomyocyte differentiations). **h,** Bystander editing efficiencies of analyzed HTS data in **g** from gDNA of R943X hiPSC-CMs seven days after eNme2-C-ABE8e treatment. The target sequence and target adenine of the most efficient sgRNA2 is shown (top). Target adenine (red) and potential bystander edits with corresponding amino acid substitutions are indicated. **i,** Representative immunofluorescence images of eNme2-C-ABE8e-sgRNA2-edited R943X hiPSC-CMs compared with Corr and R943X controls. Cells were stained for DAPI (white), MYBPC3 (magenta), and ACTN2 (cyan). Scale bar: 50 *μ*m. **j,** Schematic of the MYBPC3-containing sarcomere (left) the AlphaFold predictions (right) of stretch 867-966 in wildtype MYBPC3 (WT) and the most prevalent alleles (V944A; F942S/V944A) identified by HTS of edited R943X hiPSC-CMs. Protein structure figures were generated using PyMOL. **k,** Schematic of plasmid-based overexpression of MYBPC3 variants identified by HTS in R943X hiPSC-CMs. MYBPC3 sequences used are presented in **Table S2**. **l,** Representative images of MYBPC3 variant expression and localization. Cells were stained for DAPI (white), MYBPC3 (magenta), and ACTN2 (cyan). Scale bar: 20 *μ*m. Bars and error bars indicate mean ± s.e.m. Data in **b** were tested for normality using Shapiro-Wilk test and tested for significance using the Mann-Whitney-U test. N.D.: not detected.

Although SpCas9-derived ABEs have demonstrated high editing efficiencies across numerous disease-associated loci^32–36^, their therapeutic application to inherited cardiomyopathies is constrained by both editor size and target-site accessibility. First, the large size of SpCas9-derived base editors presents as a major translational barrier, as packaging of the editor and sgRNA cassette typically exceeds the cargo capacity of a single-AAV vector, requiring dual-AAV vector delivery strategies that limit therapeutic applicability. Second, analysis of the genomic locus surrounding the *MYBPC3* R943X variant revealed a PAM incompatible with a canonical ABE-SpCas9 sgRNA when placing the target adenine on all potential positions within the high activity window (protospacer position 4 to 8, PAM-distal base defined as position 1; **Fig. 1a**).

To overcome both size and targetability constraints, we selected eNme2-C, an engineered *Neisseria meningitidis*-derived Cas9 variant whose compact size and cytosine-rich N4CN PAM compatibility make it well suited for therapeutic gene editing applications^37–39^. Fusion of eNme2-C to the laboratory evolved adenine deaminase ABE8e generated a compact editor with an activity window spanning protospacer positions 9-16^37,39^. The consecutive cytosines in the R943X locus allowed us to design five distinct sgRNAs placing the target adenine at protospacer positions A9-A13, the region where eNme2-C-ABE8e has previously shown peak editing efficiency.

Five additional adenines fall within the editing window, posing a risk of bystander editing that could introduce unintended amino acid substitutions in MYBPC3 (**Fig. 1a**). We reasoned that ABE9, a structure-guided ABE8e variant engineered for reduced bystander editing at both adenine and cytosine positions when fused to SpCas9 orthologues, might mitigate this risk, and therefore also generated an eNme2-C-ABE9 fusion^40–42^.

To investigate the potential of compact base editors in R943X hiPSC-CMs, we constructed all-in-one lentiviral vectors encoding either eNme2-C-ABE8e or eNme2-C-ABE9, each driven by the truncated EF1α promoter (EFS) and each carrying one of the five sgRNA cassettes (**Fig. 1f**). High-throughput sequencing (HTS) of PCR-amplified genomic DNA (gDNA) one week after transduction revealed that eNme2-C-ABE8e consistently achieved the highest level of correction when combined with sgRNA2, achieving 13.05% ± 1.14% editing of the mutant allele, whereas sgRNA3 and sgRNA4 yielded substantially lower editing efficiencies (5.90% ± 0.10% and 4.0% ± 0.26%, respectively) (**Fig. 1g, h**). In contrast, we did not detect editing with any eNme2-C-ABE9 construct, despite its potent activity when paired with SpCas9 variants^40–42^. Importantly, genomic correction translated into robust restoration of MYBPC3 expression (**Fig. 1i**).

Consistent with the broad editing window of ABE8e^43,44^, correction of the pathogenic allele was accompanied by bystander editing at two coding adenines. We observed frequent bystander editing at position A6 (p.V944A, GTG>GCG; 19.39% ± 2.19%) and A12 (p.F942S, TTG>TCG; 3.64% ± 0.31%), while we observed minimal indel formation (0.63% ± 0.20%; **Fig. 1h**). Allele frequency analysis showed that most frequently edited alleles harbored the V944A substitution alone or in combination with F942S alleles, in both cases together with correction of the pathogenic target adenine (**Fig. 1h)**.

To investigate potential functional consequences of bystander edits on protein function, we interrogated publicly available human genetic variant databases, including ClinVar and gnomAD, but found no records of these specific variants^45,46^. Additionally, we inspected the effect of bystander edits on the structure of cardiac myosin filaments (PDB ID: 8G4L, **Fig. 1j**). MYBPC3 has a “beads-on-a-string“-like structure with stretches of several connected beta-sandwich domains. Stretch 867-966 contains the R943 amino acid and we predicted the structure of this stretch in the presence and absence of the ABE-induced missense variants with AlphaFold2^47^. We found that bystander edits caused no disruption to the folding of the domain *in silico*.

To assess the effect of these bystander edits on protein expression and localization, we transfected R943X hiPSC-CMs with plasmids encoding wildtype, R943X, V944A and F942S/V944A MYBPC3-variants. Three days after transfection we observed expression and sarcomeric localization of MYBPC3-variants indistinguishable from wildtype MYBPC3, indicating that the V944A and the F942S bystander substitutions do not disrupt MYBPC3 folding or incorporation into the sarcomere (**Fig. 1k, l**).

Collectively, these data establish that a compact base editor can efficiently correct the pathogenic *MYBPC3* R943X mutation and restore protein levels in patient-derived hiPSC-CMs.

### Rational engineering of a short chimeric cardiomyocyte-specific promoter

Systemically administered AAV vectors achieve widespread biodistribution and can result in substantial transgene exposure in peripheral organs, particularly the liver^48^. Although base editors are designed to minimize unintended genomic alterations through their programmable targeting, prolonged expression of the editor in non-target tissues may increase the risk of sgRNA-dependent and sgRNA-independent, genome-wide off-target editing^49–51^. As MYBPC3 is solely expressed in cardiomyocytes, restricting editor expression to this cell type could reduce unnecessary editing in non-cardiomyocytes and improve the safety profile for systemic delivery. The 1,319 amino acid size of eNme2-C-ABE8e limits promoter size to approximately 250 bp for single-AAV packaging and thereby excludes previously described short cardiomyocyte-specific promoters, such as the short *Myl2* promoter (335 bp)^52^. We therefore sought to develop a promoter that could drive cardiomyocyte-specific expression while remaining compatible with single-AAV packaging constraints. We hypothesized that promoter elements derived from genes preferentially expressed in human adult cardiomyocytes, including *MYBPC3*, *MYH6*, *MYH7*, *MYL3* and *TNNT2*, could be exploited to generate novel short chimeric promoters while maintaining cardiomyocyte specificity^53^. To do so, we computationally divided the naturally occurring 500 bp proximal promoter regions of these five genes into 50 bp fragments and reassembled these into a library of chimeric promoters of 250 bp in length for in vitro and in vivo validation (**Fig. 2a**). All five parental promoters were evenly represented within the new chimeric promoter pool (**Fig. 2b**). Novel promoter sequences were synthesized as an oligonucleotide library and inserted upstream of a GFP-poly(A) cassette in an AAV backbone. To investigate functionality of these novel chimeric promoters in hiPSC-CMs in vitro, we performed an array library screen measuring GFP expression six days after transfection (**Fig. 2c**). Immunofluorescence analysis revealed 48 chimeric promoters driving GFP expression in hiPSC-CMs (**Fig. 2d**).

**Fig. 2:**
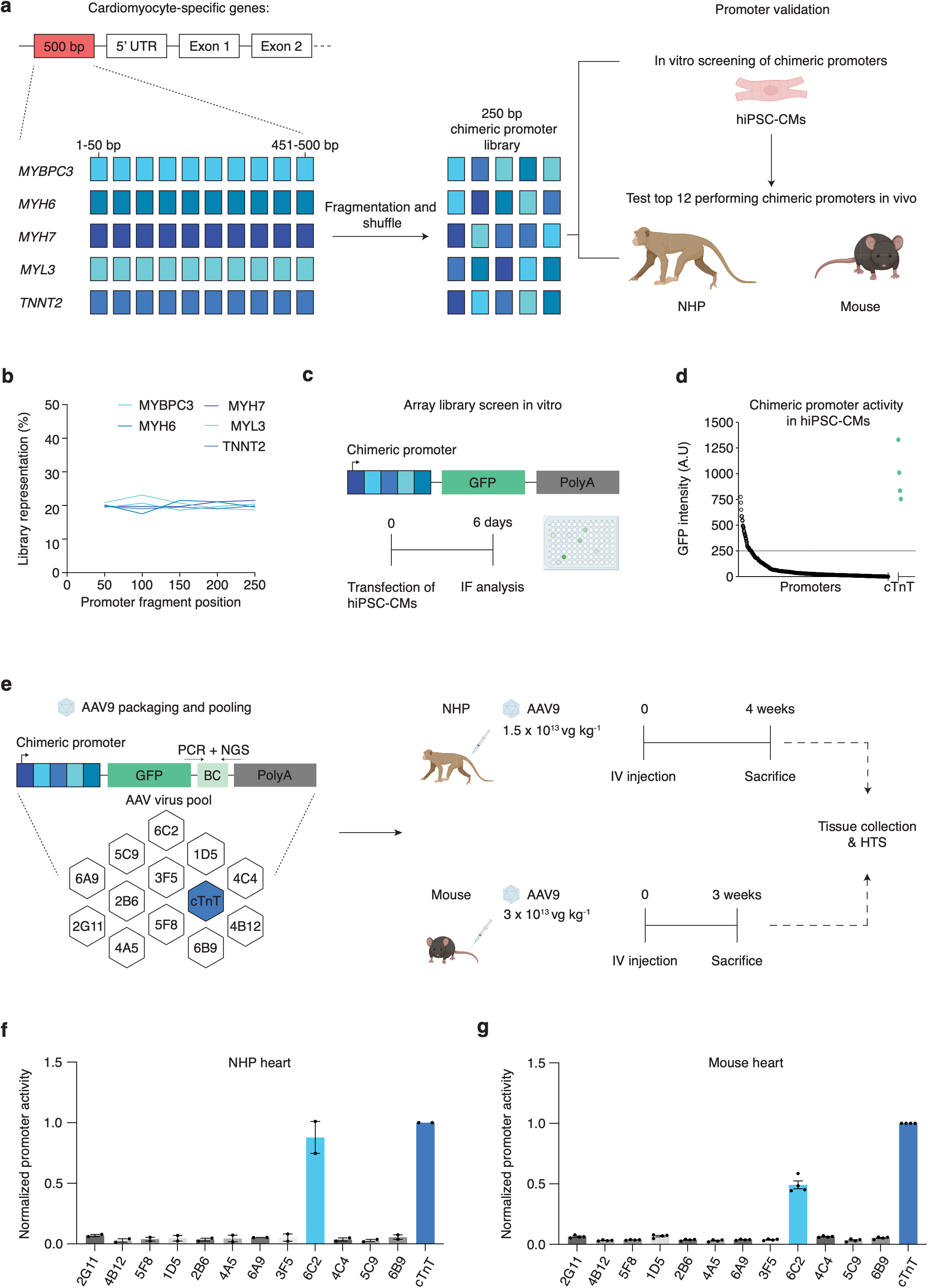
Engineering of a novel short cardiomyocyte-specific promoter in mice and non-human primates. **a**, Schematic of experimental set-up for the development of novel cardiomyocyte-specific promoters. The natural occurring 500 bp proximal promoter sequences of *MYBPC3*, *MYH6*, *MYH7*, *MYL3* and *TNNT2* were computationally digested into 50 bp fragments and randomly shuffled to generate a 250 bp chimeric promoter library *in silico*. Promoters were first screened in vitro in hiPSC-CMs and subsequently evaluated in vivo in mice and NHPs using AAV9-mediated delivery. **b**, Representation of library composition and distribution of promoter fragments present in the oligo library pool. **c**, Schematic of plasmid design and experimental set-up for the in vitro array library screen in hiPSC-CMs. **d**, Analysis of GFP intensity in hiPSC-CMs six days after transfection of plasmids containing chimeric promoters driving GFP expression. The 400 bp cTnT promoter served as positive control. **e**, Schematic of experimental set-up evaluating promoter activity in vivo in mice and NHPs. The 12 top-performing promoters from **d** were selected for in vivo validation. Promoters were packaged in AAV9 capsids driving a GFP-poly(A) cassette containing unique barcodes for identification during data analysis after HTS of gDNA and cDNA. Adult mice and NHPs were IV-injected with virus pools (3.0 × 10^13^ vg kg^−1^ and 1.5 × 10^13^ vg kg^−1^, respectively) and tissue was collected three (mice) and four (NHPs) weeks after. **f-g**, HTS analysis of PCR-amplified viral DNA and cDNA of target regions containing the barcodes in hearts of NHPs (**f**; n=2) and mice (n=4; **g**). Promoter activity was determined by comparing the relative number of barcoded transcripts versus barcoded viral DNA present in the HTS data, and normalized to activity of the 400 bp cTnT promoter. Dots represent individual mice and NHPs. Bars and error bars indicate mean ± s.e.m.

Encouraged by the high transcriptional activity of the top-performing chimeric promoters in hiPSC-CMs in vitro, we next sought to determine whether this activity translated to small- and large-animal models in vivo. Therefore, we selected the 12 top-performing candidates for evaluation in the heart and liver, the principal off-target organ for systemic delivery, of mice and NHPs (**Fig. 2e**). Each promoter was packaged with a GFP-poly(A) cassette into AAV9 capsids, a serotype that preferentially transduces cardiomyocytes compared to other cardiac cell types after systemic delivery^54^. AAVs were pooled at equal vector genome concentrations and intravenously injected at 3.0 × 10^13^ vg kg^−1^ and 1.5 × 10^13^ vg kg^−1^ in adult mice and NHPs, respectively. Heart and liver tissue was collected at three (mice) and four (NHPs) weeks post-injection followed by PCR amplification and HTS of barcoded target regions from gDNA and cDNA. Among all candidates tested, 6C2 produced strong expression in the heart, with expression levels of 0.88 ± 0.13-fold in NHPs and 0.49 ± 0.03-fold in mouse hearts compared to the cTnT promoter while maintaining minimal transcriptional activity in the liver (**Fig. 2f, g; Supplementary Fig. 1a, b**). Activity for the remaining candidates could not be confirmed in vivo.

To further characterize cardiomyocyte specificity of 6C2 in vivo, we conducted an independent mouse study comparing its transcriptional activity with that of other compact (EFS, 212 bp; U1A, 251 bp) and conventional (CMV and cTnT) promoters previously used in base editing studies ^9,55–57^. To do so, neonatal (P3) mice received a single intraperitoneal (IP) dose of 1.65 × 10^13^ vg kg^−1^ AAV9 vectors expressing eGFP (**Supplementary Fig. 2a**). Three weeks after administration, robust eGFP expression was detected in the hearts of mice injected with CMV-, cTnT- and 6C2-driven vectors, while EFS- and U1A-driven expression negligible (**Supplementary Fig. 2c**). Strikingly, eGFP expression remained undetectable in the livers of 6C2-eGFP injected mice despite a 4-fold higher AAV9 vector copy number per diploid genome (vc/dg) in the liver (10.5 ± 2.7 vc/dg) compared with the heart (2.5 ± 0.6 vc/dg), whereas cTnT-eGFP treated mice showed detectable hepatic leakage (**Supplementary Fig. 2b, d**).

Taken together, these data established 6C2 as a novel short chimeric promoter showing robust activity and specificity in cardiomyocytes of mice and NHPs in vivo.

### The 6C2 promoter drives efficient base editing in *MYBPC3* R943X mutant hiPSC-CMs

Having established 6C2 as an active cardiomyocyte-restricted promoter, we next sought to investigate its effect on base editing efficiency in hiPSC-CMs. While previous gene editing studies have shown that compact ubiquitous promoters (EFS and U1A)^55,56^ drive editor expression in mouse hearts, the extent to which promoter activity influences gene editing efficiency in cardiomyocytes remains unclear. To investigate if promoter activity influences editing efficiency, we first decided to compare 6C2 promoter activity in hiPSC-CMs with CMV, EFS, U1A, and cTnT (**Fig. 3a**)^9,57^. We expressed SpCas9-P2A-eGFP under the control of each promoter individually for seven days in hiPSC-CMs, followed by immunoblotting analysis. The CMV promoter drove strongest SpCas9 expression, followed by the 6C2 and the U1A (**Fig. 3b, c**). Surprisingly, the EFS promoter yielded minimal activity in hiPSC-CMs, despite efficient editing in previous experiments with the EFS-eNme2-C-ABE8e-sgRNA2 (**Fig. 1f-h**). This prompted us to test whether stronger editor expression driven by the 6C2 promoter could enhance editing efficiency. We therefore transduced R943X hiPSC-CMs with all-in-one lentiviral vectors encoding eNme2-C-ABE8e under the control of either the EFS or the 6C2 promoter and found substantial, but comparable proportions of MYBPC3-expressing hiPSC-CMs (**Fig. 3d**). To more quantitatively assess the percentage of cells that express MYBPC3 levels after ABE, we performed flow cytometry selecting for targeted cardiomyocytes (DAPI+, mCherry+ and TNNT2+). Editing increased only modestly with 6C2-eNme2-C-ABE8e (17.05% ± 0.99%) relative to EFS-eNme2-C-ABE8e (15.93% ± 1.37%) (**Fig. 3e, f**). This suggests that editing efficiency saturates at low effector abundance, a favorable property for single-AAV-mediated base editing strategies.

**Fig. 3:**
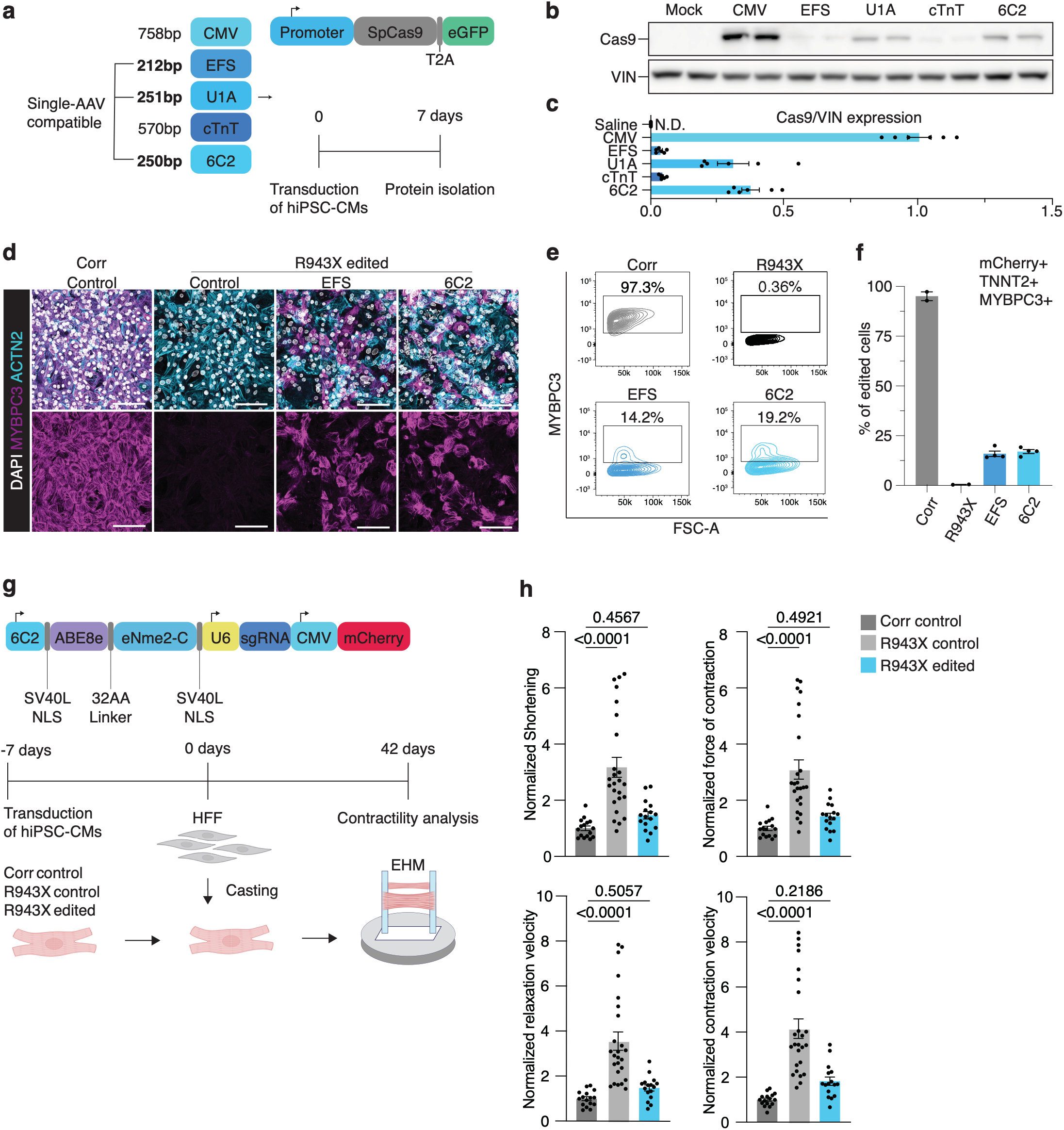
Base editing efficiency saturates at low expression levels in *MYBPC3* R943X mutant hiPSC-CMs. **a,** Schematic of promoters driving Cas9-P2A-eGFP expression used in this study. Promoters compatible with single-AAV packaging of eNme2-C-ABE8e are shown in bold. Promoter sequences are listed in **Table S3**. **b,** Western Blot of SpCas9 expression seven days after lentiviral transduction. **c,** Quantification of Western Blots from three independent cardiomyocyte differentiations (*n*=2 replicates). SpCas9 expression was normalized to VIN, GAPDH and α-tubulin. **d,** Representative immunofluorescence images of R943X hiPSC-CMs treated with EFS- or 6C2-driven eNme2-C-ABE8e and control treated R943X and Corr controls. Cells were stained for DAPI (white), MYBPC3 (magenta), and ACTN2 (cyan). Scale bar: 100 *μ*m. **e,** Representative FACS contour plots of R943X hiPSC-CMs edited with EFS- or 6C2-driven eNme2-C-ABE8e compared with Corr and R943X control-treated hiPSC-CMs. Percentages indicate MYBPC3-positive cells. MYBPC3 expression was quantified in mCherry+, TNNT2+ cells. **f,** Quantification of sorted MYBPC3-positive cells in **e**. Cells were sorted from two independent cardiomyocyte differentiations (*n*=2 controls; *n*=4 R943X-edited replicates). **g,** Schematic of EHM generation from Corr and R943X control-treated or edited R943X hiPSC-CMs. **h,** Functional analysis 42 days after casting, normalized to Corr control hiPSC-CMs (*n*=3 independent cardiomyocyte differentiations). Bars and error bars indicate mean ± s.e.m. Statistical significance was assessed by one-way ANOVA with Dunnett’s multiple comparison test. HFF: human foreskin fibroblasts; ns, not significant.

Finally, genomic editing at four computationally predicted off-target loci did not exceed background levels, aligning with the superior fidelity and heightened mismatch sensitivity of Nme2Cas9-derived base editors relative to SpCas9 orthologs (**Supplementary Fig. 3a, b**)^37,39^.

### Base editing rescues hypercontractility in *MYBPC3* R943X mutant engineered human myocardium

Loss of MYBPC3 induces hypercontractility, the same pathological hallmark that the FDA-approved myosin inhibitor mavacamten was designed to counteract in HCM patients^58–60^. We therefore asked whether R943X base editing could restore cardiomyocyte function in engineered human myocardium (EHM), a well-established 3D tissue model to study cardiomyocyte function^61,62^. hiPSC-CMs were transduced with lentiviral vectors encoding either 6C2-eNme2-C-ABE8e-control (no sgRNA) or 6C2-eNme2-C-ABE8e-sgRNA2, then combined with human fibroblasts and cast into EHM tissues for functional analysis (**Fig. 3g**). R943X EHMs displayed a hypercontractile phenotype, characterized by increased contractile force and shortened contraction and relaxation times compared to isogenic controls (**Fig. 3h**). In contrast, eNme2-C-ABE8e-edited R943X EHMs exhibited contractile force and contraction velocities comparable to those of the isogenic control tissues. Thus, in addition to restoration of MYBPC3 expression, correction of the R943X allele rescued the hypercontractile phenotype.

Together, these findings establish the all-in-one 6C2-eNme2-C-ABE8e-sgRNA2 construct as a compact, efficient platform to correct the *MYBPC3* R943X allele in hiPSC-CMs and to rescue HCM-related hypercontractility.

### Base editing of *MYBPC3* R947X mutant mouse fibroblasts

To translate an all-in-one eNme2-C-ABE8e strategy to an in vivo HCM model, we first generated a mouse line carrying the *Mybpc3* R947X variant, the murine equivalent of the human R943X allele (**Supplementary Fig. 4a**). Homozygous *Mybpc3^R^*^947^*^X/R^*^947^*^X^* (hereafter: R947X) mice showed dramatically reduced *Mybpc3* expression and lacked detectable MYBPC3 protein compared to wildtype littermates (**Supplementary Fig. 4b-e**). R947X mice developed severe cardiac hypertrophy and thickening of the ventricular walls starting from postnatal day 3 (P3) and progressing through eight weeks of age (**Supplementary Fig. 4f-l**).

Despite amino acid conservation, the murine *Mybpc3* R947X nucleotide sequence differs from the corresponding human allele at positions 5 (C>T), 13 (A>C) and 23 (C>G) within the protospacer and in the cytosine rich PAM-region (N_4_CG> N_4_AG; **Fig. 4a**). This PAM-alteration precluded translation of the human sgRNA1 with eNme2-C-ABE8e to the mouse locus, while sgRNA2-4 remained compatible. Moreover, the resulting NNNAGT PAM is compatible with the previously engineered compact SAKKH-ABE8e, thereby enabling evaluation of this additional alternative compact ABE^63^. Based on these sequence alterations, we designed four eNme2-C-ABE8e sgRNAs and one additional sgRNA for SAKKH fused to either ABE8e or ABE9 to determine the best strategy for in vivo correction. We isolated primary liver fibroblasts from two homozygous R947X mice and immortalized them with the SV40L large T antigen (**Fig. 4b**)^64^. The resulting R947X fibroblasts were transduced with all-in-one lentiviruses and subsequently sorted for mCherry to enrich for targeted cells. HTS of PCR-amplified target gDNA revealed highest on-target editing with EFS-eNme2-C-ABE8e-sgRNA4 (8.0%) followed by EFS-SAKKH-ABE8e (5.6%) with minimal indel formation (**Fig. 4c, d**). Importantly, the editing profile generated by eNme2-C-ABE8e in mouse R947X fibroblasts recapitulated the editing pattern observed in human R943X hiPSC-CMs, including the occurrence and relative distribution of bystander edits (**Fig. 4d**). In contrast, SAKKH-ABE8e displayed broader activity across targetable adenines within the protospacer, with SAKKH-ABE8e exhibiting higher overall activity than SAKKH-ABE9 (<1.0%). Based on these data, we selected eNme2-C-ABE8e-sgRNA4 and the SAKKH-ABE8e for in vivo evaluation.

**Fig. 4:**
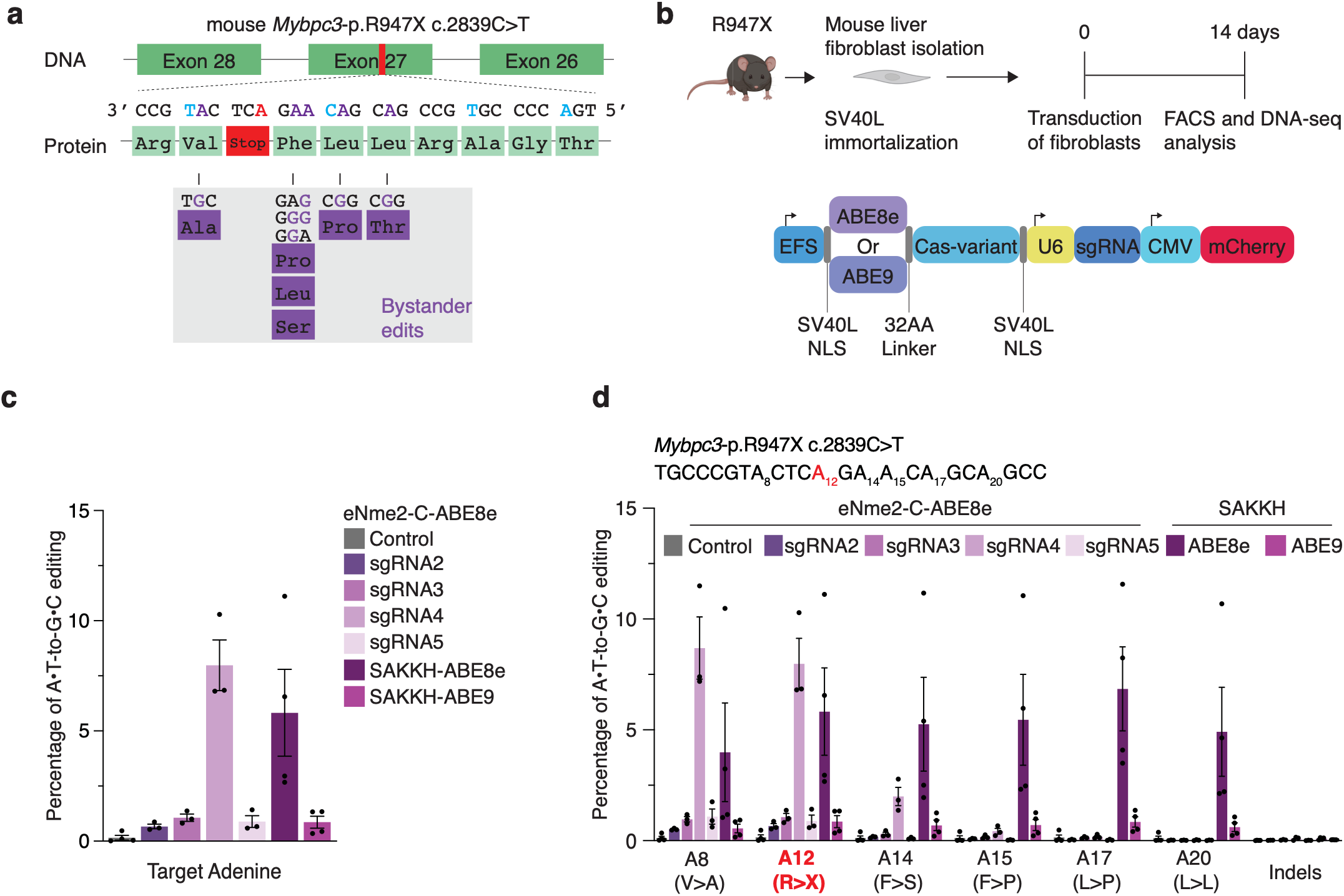
Base editing strategies to correct the *Mybpc3*-p.R947X c.2839C>T variant in mouse fibroblasts. **a,** Schematic of the R947X-containing mouse genomic region (exon 27, red). The pathogenic target adenine is shown in red. Potential ABE-induced bystander edits are shown in purple with corresponding amino acid changes. The amino acid sequence is conserved, with nucleotides differing from the human sequence shown in blue. **b,** Schematic of liver fibroblast isolation and immortalization (top) and all-in-one lentiviral constructs encoding eNme2-C or SAKKH Cas9 variants used for editing (bottom). Fourteen days after transduction, targeted fibroblasts were isolated by FACS and analyzed by HTS of PCR-amplified gDNA. **c,** HTS-quantified editing efficiency at the R947X target adenine. Data are from two independent R947X fibroblast lines (*n*=1-2 replicates per line). **d,** Bystander editing in R947X mouse fibroblasts fourteen days after transduction with base editors. The target sequence of the most efficient eNme2-C-ABE8e sgRNA4 is shown (top). Target bystander adenines and corresponding amino acid substitutions are indicated. Bars and error bars indicate mean ± s.e.m.

### Cargo-size reduction for single-AAV delivery

The stringent packaging capacity of AAV vectors (<4.7kb) necessitates extensive optimization of every component of the editing cassette to enable single vector delivery. Beyond minimizing the size of the editor and regulatory elements, compacting the sgRNA architecture represents an additional strategy to accommodate all required components within a single-AAV vector while preserving editing activity^65^. Previous work showed that the wildtype Nme2Cas9 sgRNA scaffold can be shortened from 145 to 121 nt by truncating the repeat:anti-repeat duplex without compromising nuclease activity^65^. Because the highly evolved eNme2-C-ABE8e contains numerous mutations relative to the parental Nme2Cas9-ABE8e, we first determined whether this compact sgRNA architecture remained functionally compatible with eNme2-C-ABE8e (**Supplementary Fig. 5a**). HEK293T cells harboring a lentivirally integrated *MYBPC3* R943X target sequence were co-transfected with eNme2-C-ABE8e and either the full-length or truncated sgRNA scaffold (**Supplementary Fig. 5b**). The truncated scaffold supported similar base editing efficiencies to those obtained with the wildtype scaffold, demonstrating that sgRNA minimization did not compromise eNme2-C-ABE8e activity (**Supplementary Fig. 5c**). We therefore incorporated the 121 nt sgRNA scaffold into the final vector design, reducing the total cassette size from 4713 bp to 4689 bp while maintaining robust editing activity.

### Single-AAV base editing ameliorates cardiac hypertrophy in *MYBPC3* R947X mutant mice

The major clinical and therapeutic hallmark for HCM is cardiac left ventricular hypertrophy^1^. To determine whether single-AAV-mediated ABE could therapeutically reverse cardiac hypertrophy in vivo, we packaged the size-optimized eNme2-C-ABE8e and SAKKH-ABE8e cargos into single AAV9 and AAVMYO capsids, selected for their efficient cardiac transduction and demonstrated translational potential in preclinical and clinical studies^66^. AAVMYO was of particular interest because of its enhanced cardiomyocyte targeting and improved liver de-targeting compared with AAV9, potentially enabling efficient cardiac gene editing at lower vector doses using single-AAV-mediated strategies^67,68^. Neonatal P3 R947X mice received 7.5 × 10^13^ vg kg^−1^ AAV vector genomes containing 6C2-eNme2-C-ABE8e or 6C2-SAKKH-ABE8e via single IP injection (**Fig. 5a, b**). Wildtype and R947X littermates receiving vectors lacking the locus-specific guide RNA served as controls. To evaluate whether therapeutic editing could be achieved at substantially lower vector doses using the muscle-tropic AAVMYO capsid, we injected a separate group of R947X mice with AAVMYO-6C2-eNme2-C-ABE8e at 1.5 × 10^13^ vg kg^−1^, a dose approximately 20-fold lower than most employed doses in preclinical in vivo base editing studies (typically 3 × 10^14^ vg kg^−1^)^16,17,69^. Cardiac structure and function were assessed longitudinally by echocardiography eight weeks after treatment, after which tissue was collected for molecular and histopathological analyses (**Fig. 5a, b; Supplementary Fig. 6a-c**). By eight weeks of age, control and SAKKH-ABE8e-treated R947X mice developed pronounced cardiac hypertrophy, exhibiting an approximate 2-fold increase in heart weight compared to control-treated wildtype littermates (**Fig. 5c**). In contrast, eNme2-C-ABE8e treatment (delivered by AAV9 and AAVMYO) markedly attenuated cardiac hypertrophy and left ventricular mass, restoring approximately 50% of the difference in total heart weight between R947X and wildtype mice (**Fig. 5c, d**). Echocardiographic analyses revealed 45% reduced left ventricular wall thickening towards wildtype levels during systole (**Fig. 5e, f**). Moreover, treated animals exhibited significantly reduced left ventricular volume and internal diameter (**Fig. 5g-j**). To determine whether the reduction in heart weight was accompanied by cellular remodeling, we quantified cardiomyocyte cross-sectional area (CSA). Control-treated R947X mice exhibited an increased cardiomyocyte CSA compared with wildtype littermates, consistent with cellular hypertrophy. Treatment with eNme2-C-ABE8e reduced cardiomyocyte CSA toward wildtype values, whereas SAKKH-ABE8e treatment did not improve CSA (**Fig. 5k**). Cardiomyocyte disarray is a pathological hallmark of HCM and is characterized by disruption of the highly ordered myocardial architecture leading to impaired mechanical function^70^. Histological examination by hematoxylin and eosin (H&E) staining demonstrated substantial restoration of myocardial organization in eNme2-C-ABE8e-treated R947X mice compared to control and SAKKH-ABE8e-treated littermates, indicating that molecular correction translated into structural rescue at the tissue level (**Fig. 5l**). These findings indicate that one-time administration of a compact base editor via single-AAV-mediated delivery can be sufficient to achieve therapeutic benefit in the heart.

**Fig. 5:**
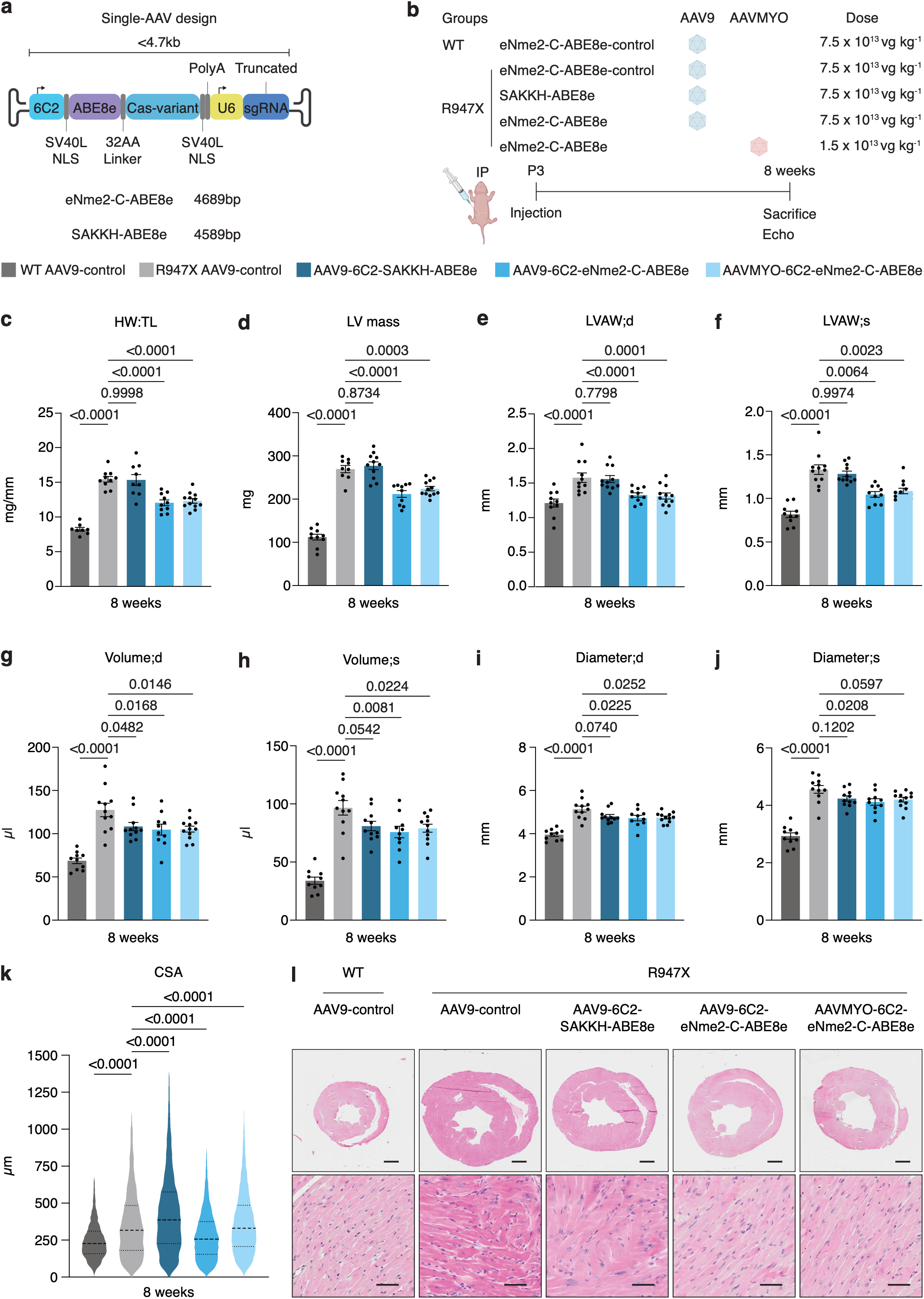
Single-AAV base editing ameliorates cardiac hypertrophy in R947X mice. **a,** Schematic of single-AAV eNme2-C-ABE8e and SAKKH-ABE8e vectors used in vivo. **b,** Experimental design for IP injections of viral vectors to P3 wildtype (*n*=10, AAV9-6C2-eNme2-C-ABE8e-control) and R947X mice (*n*=10-12 per group). **c,** Cardiac hypertrophy measured as heart weight-to-tibia length ratio (HW:TL) at eight weeks of age. **d,** Left ventricular mass after base editing. **e-j,** Left ventricular anterior wall thickness during diastole (LVAW;d) (**e**) and systole (LVAW;s) (**f**), left ventricular volume during diastole (volume;d) (**g**) and systole (volume;s) (**h**), and left ventricular diameter during diastole (diameter;d) (**i**) and systole (diameter;s) (**j**) in wildtype AAV9-control, R947X AAV9-control, and R947X-edited mice at eight weeks of age (*n*=10-12 per group). **k,** Cardiomyocyte cross-sectional area (CSA) in wildtype AAV9-control, R947X AAV9-control, and R947X-edited mice (*n*=3-5 hearts per group; *n*=2520-3250 cells per group). Sections were stained with Wheat Germ Agglutinin (WGA), and CSA was quantified using Cellpose2.0^98^. **l,** Representative H&E-stained heart sections from wildtype AAV9-control, R947X AAV9-control, and R947X-edited hearts at eight weeks of age (top). Restored cardiomyocyte disarray observed in eNme2-C-ABE8e-treated R947X mice (bottom row). Scale bars: 1000 *μ*m (top) and 50 *μ*m (bottom). Bars and error bars indicate mean ± s.e.m. Statistical significance was assessed by ordinary one-way ANOVA and Dunnett’s multiple comparison test (**c-j**) or by the Kruskal-Wallis test (**k**); ns, not significant.

### Molecular assessment of single-AAV-mediated ABE in vivo

To determine the efficiency of in vivo base editing at gDNA in the heart, we quantified on-target editing eight weeks after neonatal AAV administration. Among all treatment groups, AAV9-eNme2-C-ABE8e achieved the highest genomic editing in ventricular tissue, with a mean editing efficiency of 10.87% ± 1.31%, followed by AAVMYO-eNme2-C-ABE8e (6.99% ± 2.18%) and AAV9-SAKKH-ABE8e (0.17% ± 0.03%) (**Fig. 6a**). Because base editor expression is restricted to cardiomyocytes, which comprise approximately 30% of the cells in the adult mouse heart^71^, bulk genomic editing frequencies underestimate the true editing rate within the therapeutically relevant cardiomyocyte population. We therefore quantified editing at the transcript level and found that AAV9-eNme2-C-ABE8e corrected 46.34% ± 2.75% of *Mybpc3* transcripts on average, with individual animals reaching 59.94% (**Fig. 6b**), confirming efficient editing of cardiomyocytes in vivo. We next assessed whether genomic correction of the mutant allele restored endogenous *Mybpc3* expression levels. AAV9-eNme2-C-ABE8e treatment restored *Mybpc3* mRNA expression to approximately 40% of wildtype levels (**Fig. 6c**). Restoration of transcript expression translated into recovery of MYBPC3, as confirmed by Western Blot analysis of both AAV9- and AAVMYO-eNme2-C-ABE8e-treated R947X hearts, with protein expression reaching up to 64.2% of wildtype levels (**Fig. 6d**). Immunofluorescence staining further demonstrated widespread restoration of MYBPC3 expression throughout the heart in ventricular cardiomyocytes following treatment with either a high or low dose of respectively AAV9- and AAVMYO-mediated delivery of eNme2-C-ABE8e (**Fig. 6e**).

**Fig. 6:**
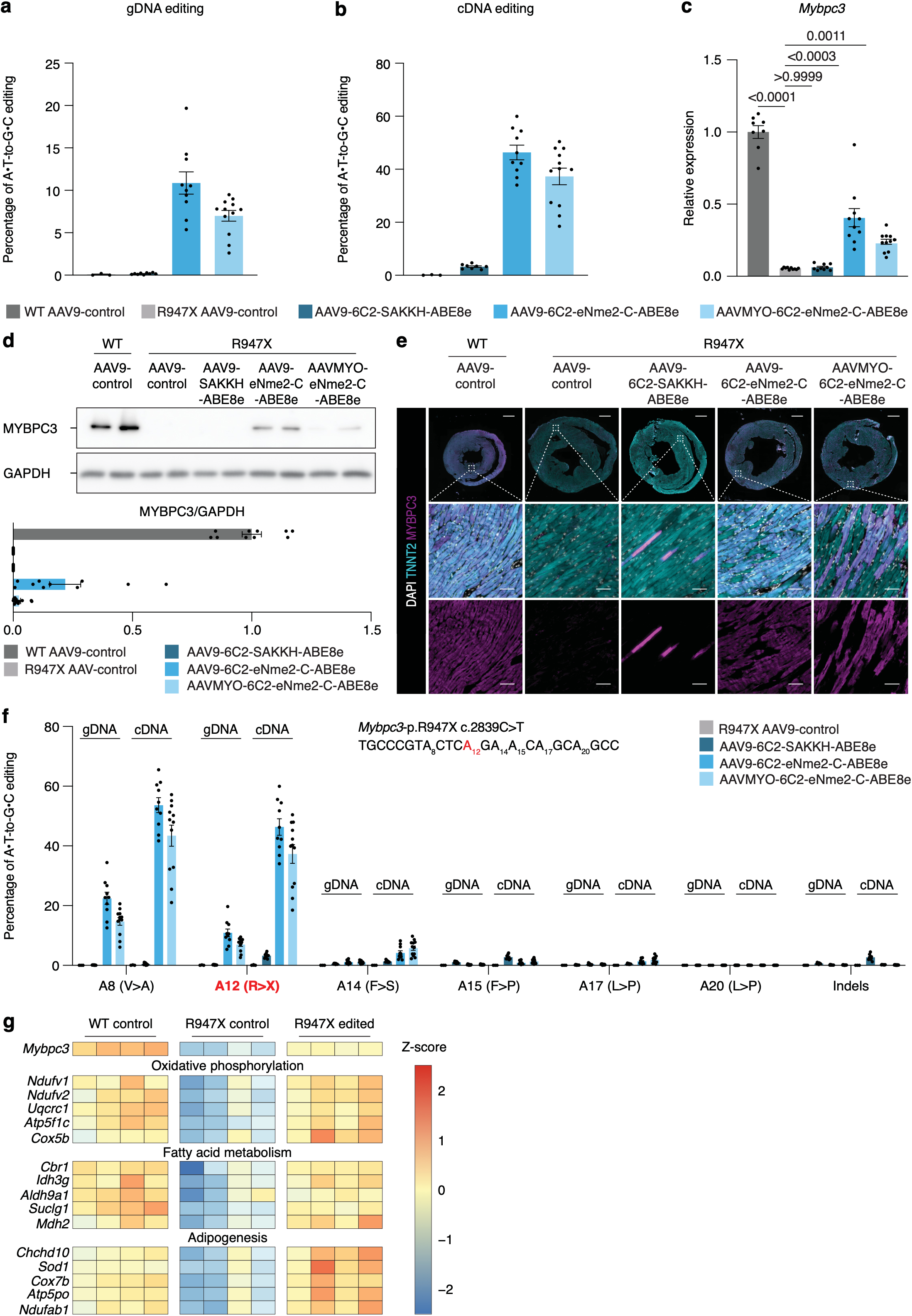
Single-AAV-mediated base editing efficiently reverts the pathogenic R947X mutation and restores protein expression in mouse hearts in vivo. **a,** Target adenine editing efficiency on gDNA in control and ABE-treated mouse hearts eight weeks after a single injection (*n*=3 R947X AAV9-control; *n*=10-12 R947X-edited). **b,** Editing efficiency in PCR-amplified cDNA from edited and control mouse hearts (*n*=3 R947X AAV9-control; *n*=10-12 R947X-edited). **c,** qPCR analysis of *Mybpc3* expression in wildtype AAV9-control (*n*=8), R947X AAV9-control (*n*=10), and R947X-edited (*n*=9-12) mouse hearts. Primers are listed in **Table S1**. **d,** Representative MYBPC3 Western Blot after eNme2-C-ABE8e editing (top) with quantification (bottom). MYBPC3 expression was normalized to GAPDH (*n*=9-12 per group). **e,** Representative immunofluorescence images of transverse heart sections from wildtype AAV9-control, R947X AAV9-control, and R947X-edited mice. Hearts were stained for DAPI (white), TNNT2 (cyan), and MYBPC3 (magenta). Scale bars: 1000 *μ*m (top) and 50 *μ*m (middle and bottom). **f,** Bystander editing analysis in PCR- amplified gDNA and cDNA from edited R947X mouse hearts. The target sequence of the most efficient eNme2-C-ABE8e sgRNA4 is shown (top middle). Target adenines and corresponding amino acid substitutions are indicated. **g,** RNA-seq heatmaps of genes restored by ABE in R947X mice relative to wildtype mice (*n*=4 per group). Bars and error bars indicate mean ± s.e.m. (**a-d, f**).

Given the bystander editing observed in edited human cardiomyocytes, we next evaluated editing outcomes in treated mouse hearts. Consistent with our in vitro findings, the predominant bystander event was the V>A substitution (22.46% ± 2.03%), whereas editing of the F>S bystander site remained minimal (1.10% ± 0.19%; **Fig. 6f**). This increase in V>A and reduction in F>S editing in mice likely reflects the two-nucleotide shift in the murine protospacer alignment, which repositions the corresponding adenine relative to the high-activity center of the eNme2-C-ABE8e editing window.

A key challenge for the clinical translation of systemically delivered AAV base editors is achieving efficient on-target editing in therapeutically relevant tissue, while minimizing unintended on-target and off-target editing elsewhere, such as genome-wide DNA and transcriptome-wide RNA editing. To evaluate tissue specificity in vivo, we collected the heart, liver, lung, diaphragm, gastrocnemius muscle, spleen and kidney from control- and eNme2-C-ABE8e-sgRNA4 treated mice and quantified vector biodistribution alongside on- and off-target genomic editing by HTS. As expected, AAV9 exhibited broad systemic biodistribution, with the highest vector copy numbers detected in the diaphragm, followed by the liver and heart (**Supplementary Fig. 7a**). In line with previous reports, AAVMYO efficiently transduced diaphragm, heart and skeletal muscle^67,72^. Notably, we observed a relative reduction of liver transduction, consistent with the reported liver de-targeting properties of muscle-tropic AAV capsids^67^. Despite efficient vector delivery to several peripheral tissues, we detected no on-target editing, indel formation or off-target editing above background levels of control-injected mice in the lung, diaphragm, gastrocnemius muscle, spleen or kidney (**Supplementary Fig. 7b, c; Supplementary Fig. 8a, b**). Low but detectable on-target editing was observed only in the liver (±2%), whereas robust editing remained restricted to the heart. Notably, no detectable editing was observed in the diaphragm or gastrocnemius despite high vector copy numbers in both tissues (69.77 ± 16.93 and 19.45 ± 4.86 vc/dg, respectively). Considering that robust editing was achieved in vitro even with the weak EFS promoter, this absence of editing in highly transduced peripheral tissues strongly suggest that the 6C2 promoter drives negligible editor expression outside the heart, functionally restricting base editing to cardiomyocytes.

Finally, to identify the molecular consequences of *Mybpc3* R947X base editing more deeply, we performed transcriptome-wide RNA-sequencing (RNA-seq) of wildtype, R947X control, and R947X AAVMYO-eNme2-C-ABE8e treated hearts. RNA-seq analysis revealed *Mybpc3* as one of the top differentially expressed genes after editing and pathway analysis revealed restored gene expression related to oxidative phosphorylation, fatty acid metabolism and adipogenesis, all patterns of metabolic normalization consistent with previous dual-AAV base edited mouse models of sarcomeric HCM (**Fig. 6g; Supplementary Fig. 9a, b**)^18,73^.

Overall, these data demonstrate the therapeutic potential of single-AAV, cardiomyocyte-restricted base editing for correcting a clinically relevant *Mybpc3* nonsense variant in vivo, achieving durable molecular correction and reversal of cardiac hypertrophy at a fraction of the dose being used in clinical trials. Importantly, these findings show the potential to modify disease progression and ultimately improve the lives of patients suffering from inherited cardiomyopathies.

## Discussion

While base editing has successfully entered the clinics for targeting the liver using lipid-nanoparticle (LNP) based delivery of the editor components^74,75^, efficient LNP-mediated targeting of cardiomyocytes via systemic injection has so far not been established^76,77^. AAV-mediated delivery is promising, but the clinical translation of earlier dual-AAV base editing strategies is limited by the high viral titers required for therapeutic efficacy^16–18,78^, as well as the limited capacity of single-AAV vectors to combine compact ABEs with tissue-specific promoters to restrict on- and off-target editing in non-target tissues^56,79–82^. Here, we tackle both limitations by co-packaging compact base editors with our newly developed minimal 6C2 promoter into single cardiotropic AAVs. By using a next generation capsid with enhanced cardiac targeting and liver de-targeting properties, we achieved therapeutic benefit in the heart at doses 20-fold lower than state-of-the-art cardiomyopathy base editing studies^16,17,66,83^, while minimizing on- and off-target editing in non-cardiac tissues. Although further studies are needed to assess safety and long-term efficacy, we anticipate that next generation single-AAV-mediated strategies are well positioned to replace dual-AAV approaches to improve the prospects for clinical translation in patients with genetic disease.

While we and others have achieved therapeutic benefit in the murine heart, the observed potentially harmful bystander editing might constrain first-in-human clinical base editing trials for cardiomyopathy patients^18,66,73,78,84^. Extensive engineering of highly active deaminase domains has come at the cost of broad editing windows, consistent with the high bystander rates we observed using ABE8e. We reasoned that ABE9 could offer a direct route to reduced bystander activity, since it was engineered specifically to achieve higher precision without loss of efficiency^40^. Contrary to this expectation, fusing ABE9 to either eNme2-C or SAKKH yielded little to no detectable activity. These observations align with a recent independent, systematic characterization of Nme2Cas9-derived base editors, in which ABE9-Nme2Cas9 fusions failed to exceed background editing^39^. TadA-9e-Nme2Cas9 variants in genomic regions with a compatible PAM might offer therapeutic utility for gene editing applications where bystander editing should be limited. Taken together, these findings suggest that deaminase properties cannot necessarily be transferred directly from SpCas9-based editors to compact Cas9 orthologues. Instead, deaminase engineering should be performed in the context of the intended Cas9. Further development of deaminase variants specifically optimized for compact Cas9 proteins may therefore enable the generation of smaller and more precise base editors with improved safety profiles for patients.

Non-cardiac base editor expression could limit therapeutic application by the generation of sgRNA-dependent or -independent off-target editing in undesired tissues. While cardiomyocyte-specific ABE has been achieved in dual-AAV approaches by utilizing the well-established cTnT promoter, single-AAV strategies have heavily relied on small ubiquitous promoters due to packaging constraints. In our study, we developed the 250 bp chimeric cardiomyocyte-specific 6C2 promoter and achieved robust cardiomyocyte-specific editing with negligible editing in other AAV targeted organs. Combining the cardiomyocyte-specificity of the 6C2 promoter with the cardiac tropism of AAV9 and AAVMYO capsids^54,67^ likely enhanced editing specificity. Importantly, our initial finding that even minimal base editor expression is sufficient to reach an editing plateau suggests that the 6C2 promoter maintains transcription strictly below this functional threshold in off-target tissues, demonstrating exceptional cardiomyocyte specificity. Future studies will provide more knowledge whether this cardiomyocyte-specificity observed in mice and NHPs will translate to the human heart. A recent report has demonstrated robust cardiomyocyte-specific gene silencing of Camk2d in mouse hearts using a 4951 bp expression cassette containing SauriCas9-ABE8e driven by the *Myl2* promoter^52^. Additionally, high-throughput evaluation of cardiomyocyte-specific promoters revealed considerable *Myl2* promoter activity in the liver in vivo^85^. Building on these findings, we incorporated the 250 bp 6C2 promoter, a minimal poly(A) signal and a truncated sgRNA scaffold reducing our AAV cargo size to a final 4689 bp, significantly advancing feasibility for large clinical-scale manufacturing^7,19,86^.

By restricting ABE expression via cardiotropic vectors and cardiomyocyte-specific promoters, actual editing efficiency within target cardiomyocytes is likely to be substantially higher than indicated by bulk tissue analysis^71^. Prior work has shown that dual-AAV SpRY-ABE8e-mediated A>G correction of 9.56% in bulk cardiac tissue was sufficient to outperform MYBPC3 replacement therapy and revert cardiac structure and function back to wildtype levels^18^. Interestingly, in our study we achieved higher average A>G conversion (10.9%) with lower therapeutic doses, while we observed only partial structural and functional recovery. Importantly, both approaches displayed similar bystander editing patterns, indicating that these secondary edits might not compromise therapeutic efficacy or cardiomyocyte viability. One key explanation for this discrepancy lies in the timing of therapeutic intervention. We administered ABEs at P3 when pathological cardiac hypertrophy was already established in R947X mice, while in the prior study, vectors were administered at P0 enabling gene correction before disease onset and corrected cardiomyocytes were still proliferating.

Intervention after disease was shown to limit the extent of phenotypic rescue, despite full restoration of MYBPC3 protein expression after gene replacement therapy^10^. Nevertheless, our findings demonstrate potentially broad clinical application since base editing after disease onset can halt disease progression and partially reverse established cardiac hypertrophy.

To translate base editing into the clinics it is essential to carefully assess the therapeutic threshold in target tissues for treating genetic diseases. Here, we demonstrate that ABE effectively restores MYBPC3 expression in hiPSC-CMs with subsequent restoration of cardiomyocyte function measured using EHMs. Building on prior preclinical studies demonstrating phenotypical rescue in hiPSC-CMs^16,72,87^, we sought to extend these findings by applying base editing directly to differentiated cardiomyocytes rather than targeting undifferentiated hiPSCs or expanded single-edited clones. We achieve up to ±20% ABE-mediated genetic correction in differentiated cardiomyocytes, demonstrating that a mosaic population of hiPSC-CMs harboring variable editing outcomes is sufficient to yield robust functional rescue of HCM-related hypercontractility^59,60^. In addition, we provide evidence that a single dose of single-AAV-mediated ABE can achieve therapeutic benefit in a cardiomyopathy-related disease model in vivo. We show that 5-20% correction in bulk cardiac tissue can reduce 50% of the disease-associated hypertrophy. Prior studies demonstrated that ±10% editing in bulk ventricular tissue can achieve near complete rescue of the HCM phenotype in mice^16–18^. Moreover, that partial correction can achieve therapeutic benefit is supported by interim data of the MyPEAK-1 HCM trial for MYBPC3 gene replacement therapy, where an average 4% increase in MYBPC3 protein led to reduced ventricular mass index and hypertrophy, and improved patients to NYHA Class I^11^. The reported low biological thresholds are of crucial clinical significance, because it would indicate that base editing platforms can prioritize precision and fidelity over maximal efficiency. However, whether the modest protein increase reported in the MyPEAK-1 trial reflects uniform, low-level expression across the entire myocardium or robust expression concentrated within a subpopulation of transduced cardiomyocytes remains unknown. In future preclinical and clinical base editing studies it is important to study the therapeutic threshold in a disease specific setting.

As next steps towards clinical translation, long-term evaluation of the potential functional consequences of bystander edits needs to be assessed. Additionally, given that 3D chromatin organization dictates ABE genomic accessibility^52,88^, off-target profiling must be performed in clinically relevant human cell-types. Advanced translational models with mature cardiac phenotypes, such as human cardiac slices from myectomy tissue from HCM patients, better represent clinically relevant profiles^89–92^. Additionally, given the potential for unanticipated editing events, complementary unbiased genome-wide approaches may provide a more comprehensive assessment of off-target activity in the context of therapeutic development^93,94^. Although we show a therapeutic benefit of ABE-mediated correction after injection in pups, efficacy studies in large animal models will further help define the therapeutic thresholds, dose selection and determine potential safety issues of AAV-mediated toxicities. Moreover, the incomplete penetrance of *MYBPC3*-associated HCM makes it difficult to predict who to treat and at which time-point during life for disease prevention or curation^95,96^. As last, despite us showing partial reversal of cardiac hypertrophy in mice, it remains unclear whether genetic correction could reverse the reported fibrosis in HCM patients^97^.

In conclusion, our findings demonstrate potential reversal of genetic heart disease by using compact tissue-restricted base editors with negligible off-target activity. We anticipate that single-AAV-mediated tissue-specific base editing strategies will enhance therapeutic safety, streamline clinical translation and ultimately can lead to a potential cure that dramatically improves the lives of patients suffering from genetic disease.

---

**Supplementary Fig. 1:**
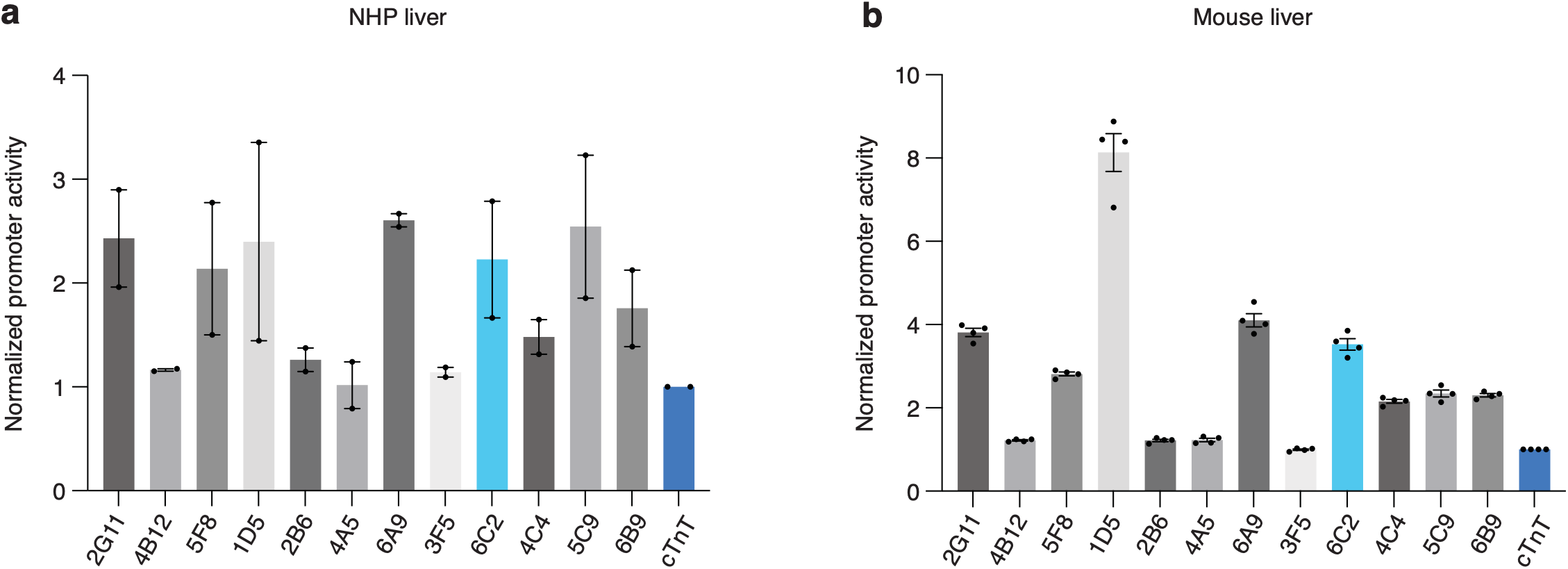
Chimeric promoter activity in the heart and liver of mice and non-human primates. Analysis of promoter activity using HTS of PCR-amplified viral DNA and cDNA of barcoded target regions in livers of NHPs (**a**; n=2) and mice (**b**; n=4). Relative promoter strengths were determined by comparing the relative number of barcoded transcripts versus barcoded viral DNA present in the HTS data. The 400 bp cTnT promoter was used as positive control. Dots represent individual mice and NHPs. Bars and error bars indicate mean ± s.e.m.

**Supplementary Fig. 2:**
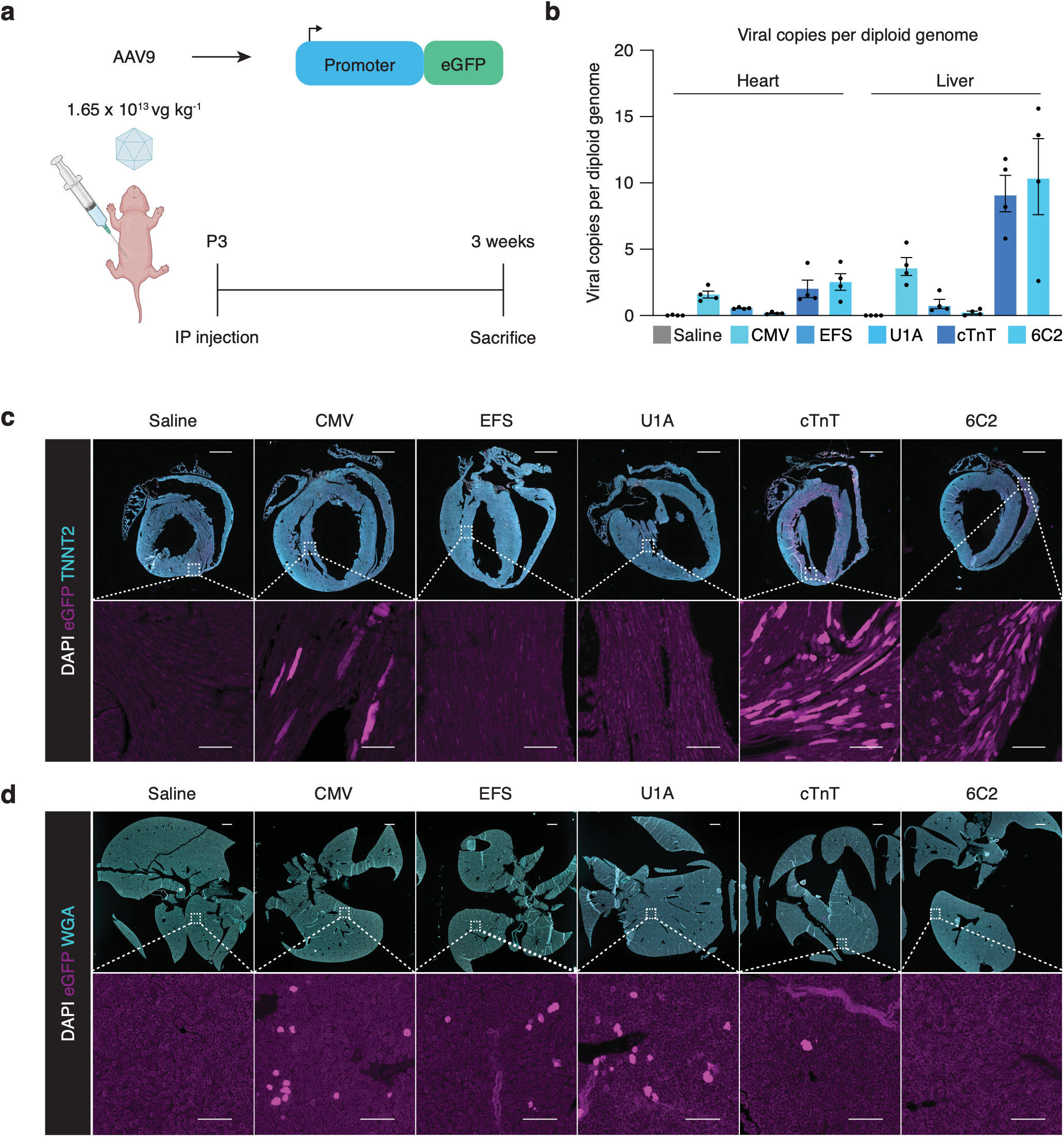
The 6C2 promoter shows stringent cardiomyocyte-specific expression in vivo. **a,** Schematic of injections with 1.65 × 10^13^ vg kg^−1^ AAV9 vector genomes containing different promoters driving eGFP expression. Mice were injected at P3 and sacrificed at 3 weeks of age. Mice injected with saline served as control. **b,** Viral copy number per diploid genome (vc/dg) in heart and liver tissues (*n*=4 per group) of mice treated with AAV9 containing the different promoters. The primers and probe bind to an independent U6 promoter upstream of the small promoters driving eGFP expression (**Table S4**). The AAV9 viral copy number per diploid genome was normalized to the mouse glucagon gene copy number in the same assay. **c,** Immunofluorescence images of heart sections (top row) and zoom (bottom row) of hearts from mice treated as depicted in **a**. DAPI (white), TNNT2 (cyan) and eGFP (magenta). Scale bar: 1000 *μ*m (top) and 100 *μ*m (bottom). **d,** Immunofluorescent images of liver sections (top row) and zoom (below) from treated mice in **c**. DAPI (white), WGA (green) and eGFP (magenta). Scale bar: 1000 *μ*m (top row) and 100 *μ*m (bottom row). Dots represent individual mice in **b**. Bars indicate mean ± s.e.m. Promoter sequences are presented in **Table S3**.

**Supplementary Fig. 3:**
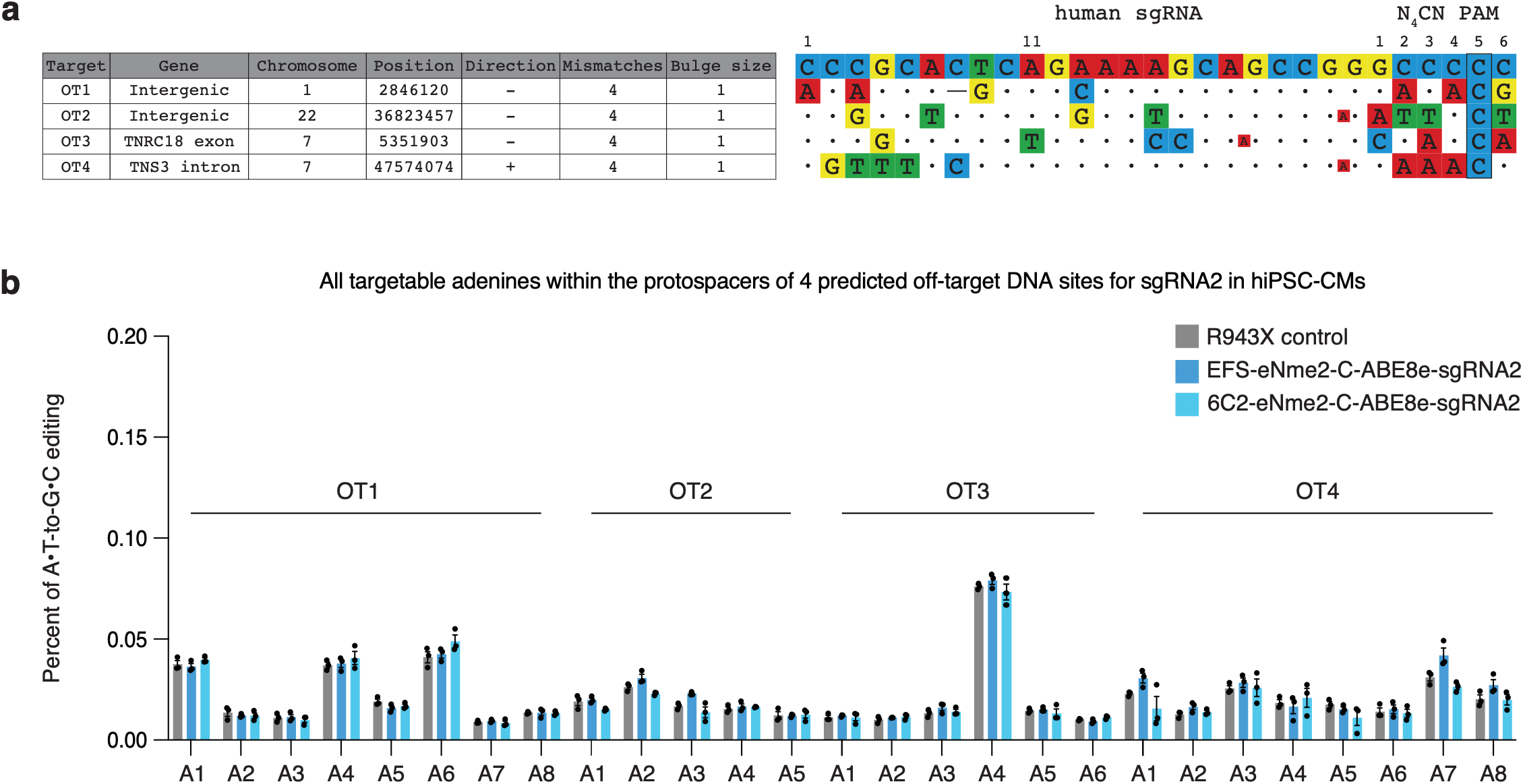
Computationally predicted sgRNA-dependent off-targets for eNme2-C-ABE8e-sgRNA2. **a,** Table (left) and sequences (right) of four Cas-OFFinder-predicted candidate off-target (OT) sites. sgRNA-matching nucleotides are shown as dots and mismatches as colored nucleotides. RNA bulges are indicated by vertical lines and DNA bulges by additional nucleotides within the protospacer. Protospacer numbering begins at the PAM-distal 5′ nucleotide (position 1). **b,** HTS-quantified editing efficiency across all 28 targetable adenines within the protospacers of four predicted candidate OT loci in control-treated and eNme2-C-ABE8e-sgRNA2-treated hiPSC-CMs driven by the EFS or 6C2 promoter (*n*=3 independent cardiomyocyte differentiations). HTS data were analyzed using CRISPResso2 and are presented as raw percentages of sequencing reads. PAM, protospacer adjacent motif; N, A, T, G, or C; OT, off-target. Bars and error bars indicate mean ± s.e.m.

**Supplementary Fig. 4:**
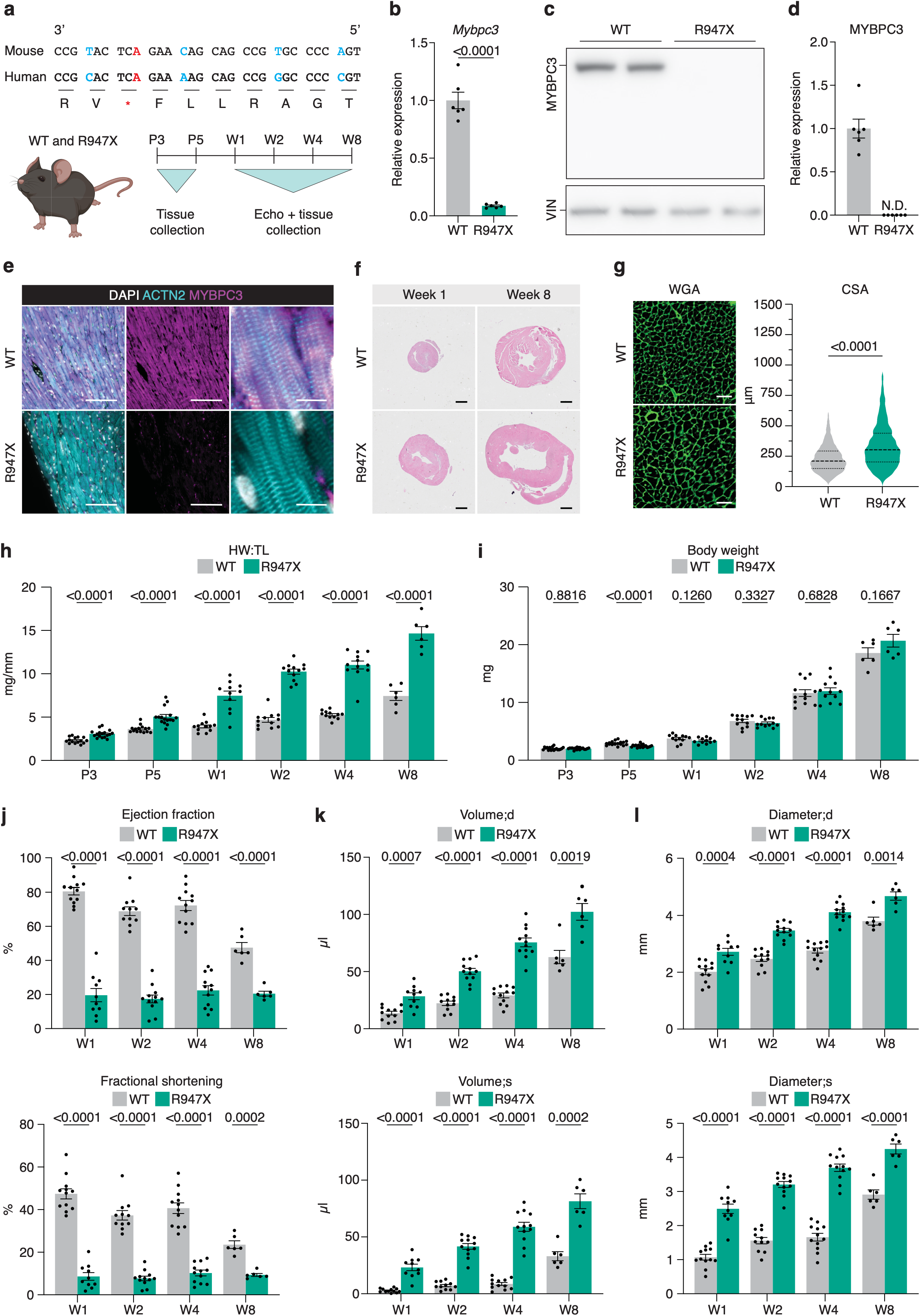
Homozygous R947X mice present with severe cardiac hypertrophy and heart failure. **a,** Schematic of the genomic region of the mouse equivalent (R947X) of the human pathogenic R943X mutation (top) and the study design (bottom). The amino acid sequence is conserved. Silent DNA nucleotide substitutions are depicted in blue. The pathogenic adenine is highlighted in red. **b,** Analysis of *Mybpc3* expression by qPCR in wildtype and R947X mice (*n*=6 for each genotype). Expression levels were normalized to *Hprt*, *Rpl32* and *18s*. **c,** Representative Western Blot of MYBPC3 expression in wildtype and R947X mice. **d,** Quantification of MYBPC3 expression in wildtype and R947X mice. MYBPC3 expression was normalized to VIN. **e,** Representative immunofluorescence staining of MYBPC3 expression and localization in wildtype and R947X mice hearts (DAPI, white; ACTN2, cyan; MYBPC3, magenta). Scale bars: 100 *μ*m (left and middle) and 10 *μ*m (right). **f,** Representative H&E staining of transverse heart sections from wildtype and R947X mice at the age of one (left) and eight (right) weeks of age. Scale bar: 1000 *μ*m. **g,** Representative WGA (left) and quantification (right) of cross-sectional area (CSA) of cardiomyocytes from wildtype and R947X mice at 8 weeks of age (*n*=3 mice per genotype; *n*=2118-2705 cardiomyocytes per group). Statistical significance was assessed by the Mann-Whitney-U test. Scale bar: 50 *μ*m. **h,** Quantification of heart size by measuring heart weight relative to tibia length in wildtype and R947X mice at all time points. **i,** Body weights of wildtype and R947X mice at the depicted time points in **a**. **j,** Analysis of cardiac function by measuring ejection fraction (top) and fractional shortening (below) from wildtype and R947X mice at one, two, four and eight weeks of age. **k-l,** Left ventricular volume (**k**) and diameter (**l**) during diastole (d; top) and systole (s; bottom) in wildtype and R947X mice at different time points. Individual dots represent individual mice. Bars and error bars indicate mean ± s.e.m. For **h-l** *n*=6-18 mice per genotype. Statistical significance between two groups was assessed by a Welch’s t-test.

**Supplementary Fig. 5:**
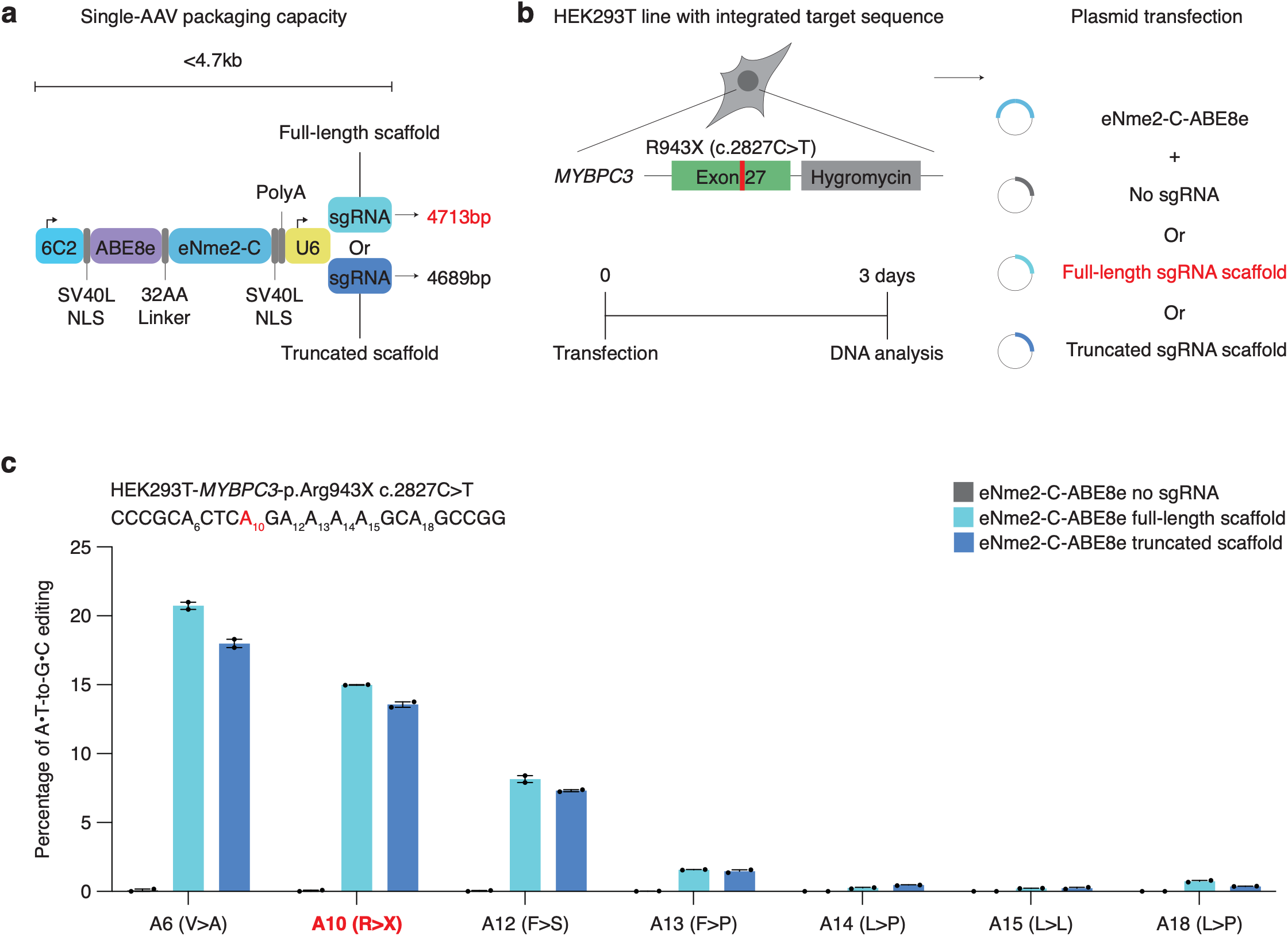
Assessment of sgRNA scaffold architecture on ABE efficiency for eNme2-C-ABE8e in HEK293T cells. **a,** Schematic of the all-in-one AAV vector design when combined with the full-length (145 nt) or the truncated (121 nt) scaffold variants that have been developed for Nme2Cas9 variants^65^. The 4713 bp construct exceeds the AAV practical packaging capacity (red). **b,** Schematic of a HEK293T cell line harboring the randomly integrated R943X-hygromycin target sequence via lentiviral transduction followed by antibiotic selection. **c,** HTS analysis results of PCR-amplified gDNA of the integrated R943X target sequence three days after co-transfection with eNme2-C-ABE8e and the full-length (*n*=2) or truncated (*n*=2) sgRNA2. Cells transfected without the sgRNA served as controls (*n*=2). Bars and error bars indicate mean ± s.e.m. Scaffold sequences are depicted in **Table S4**.

**Supplementary Fig. 6:**
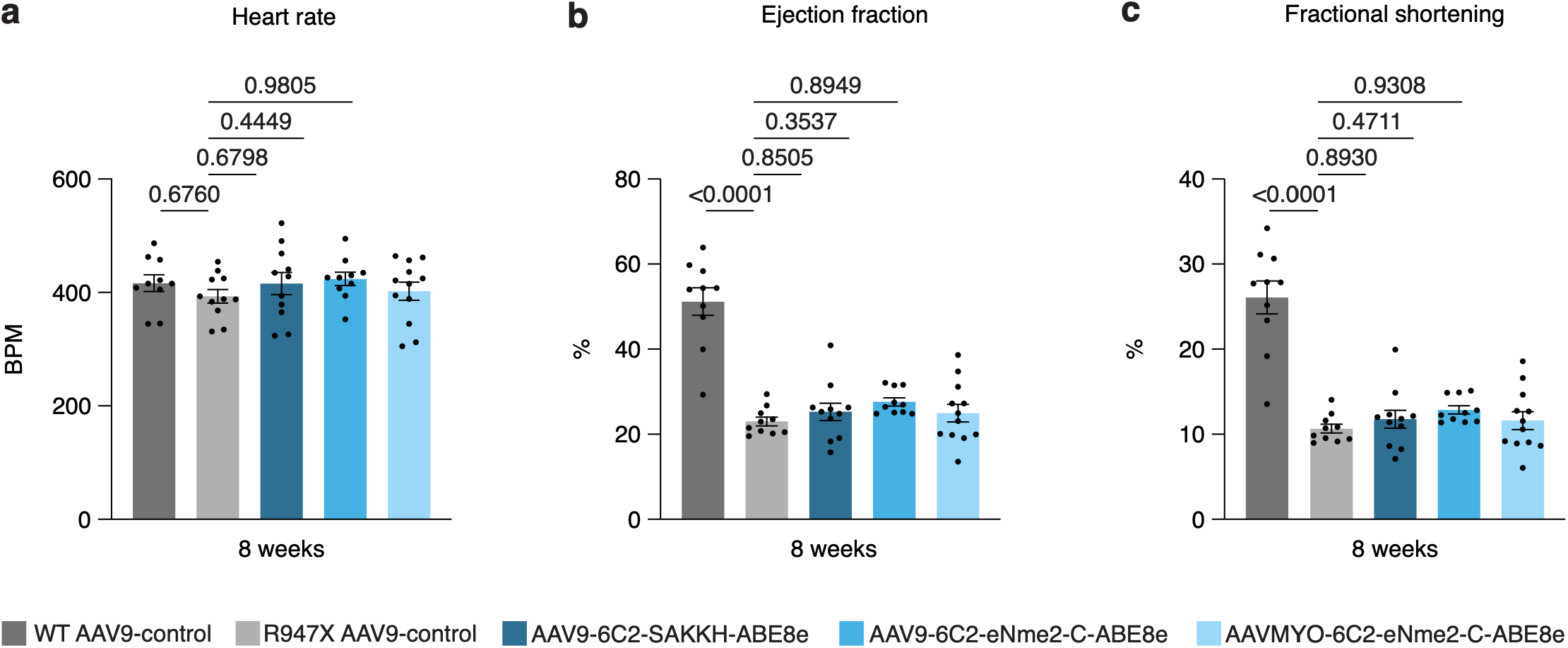
Base editing ameliorates cardiac dysfunction in vivo. Heart rate **a**, EF **b**, and FS **c,** of wildtype AAV9-control, R947X-control and R947X edited mice at 8 weeks of age (*n*=10, wildtype AAV9-control; R947X AAV9-control and R947X-edited mice (*n*=10-12 per group). Control AAV9 contained 6C2-eNme2-C-ABE8e without the sgRNA sequence. Bars and error bars indicate mean ± s.e.m. Statistical significance was assessed by one-way ANOVA and Dunnett’s multiple comparison test.

**Supplementary Fig. 7:**
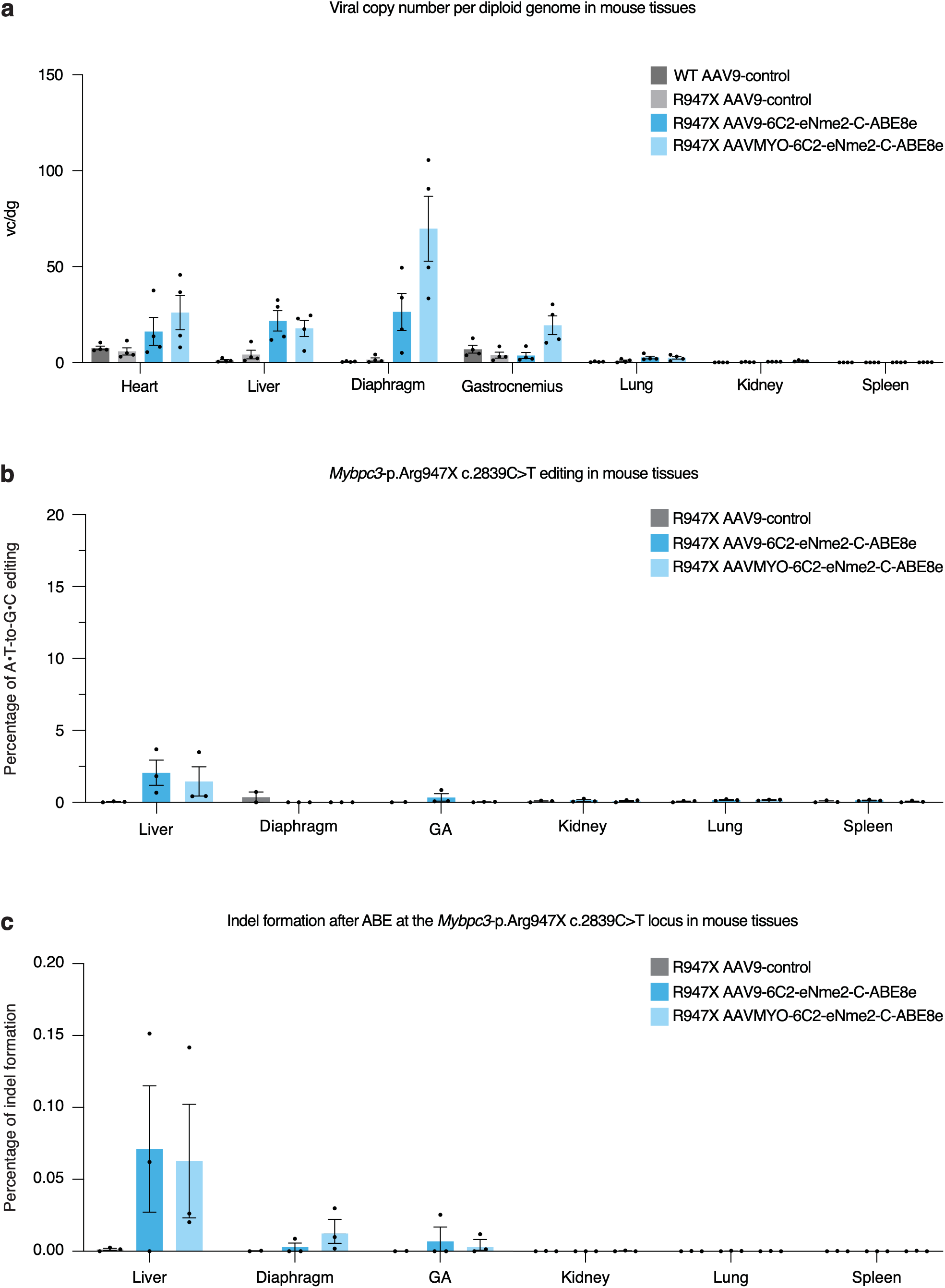
AAV vector biodistribution, on-target ABE and indel formation in mouse tissues eight weeks after treatment with single-AAV ABEs. **a,** Mice were injected at P3 with 7.5 × 10^13^ vg kg^−1^ AAV9 and 1.5 × 10^13^ vg kg^−1^ AAVMYO vector genomes (*n*=4 per group) and sacrificed at eight weeks of age. Primers and probe anneal to the 6C2 promoter of the transgene in both AAV9 and AAVMYO (**Table S4**). The copy number of the mouse glucagon gene was used to normalize to the AAV copy number per diploid genome^99^. **b**, Analysis of HTS data to measure on-target ABE efficiency in the liver, diaphragm, gastrocnemius, kidney, lung and spleen. HTS was performed on PCR-amplified gDNA of R947X AAV9-control mice and R947X eNme2-C-ABE8e edited mice delivered by AAV9 or AAVMYO (*n*=2 or 3 per group). **c**, Indel formation after ABE of the same samples analyzed in **b**. Dots represent individual mice. Bars and error bars indicate mean ± s.e.m.

**Supplementary Fig. 8:**
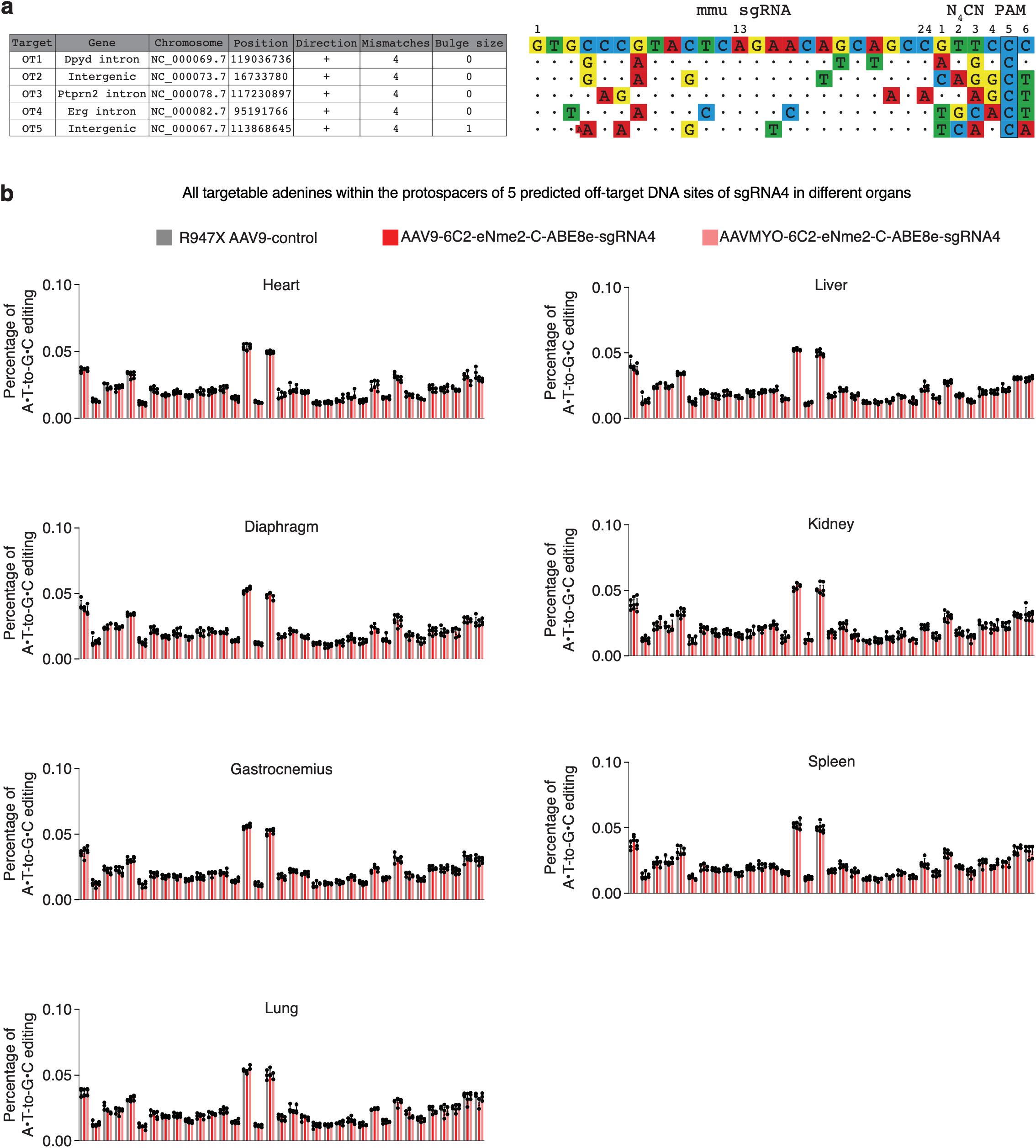
Off-target editing at computationally predicted sgRNA-dependent off-target sites of the mouse R947X locus for sgRNA4 combined with eNme2-C-ABE8e. **a,** Table (left) and sequence (right) depicting five candidate off-target (OT) sites predicted using Cas-OFFinder^100^. Nucleotides that match the sgRNA sequence are depicted as dots. Mismatches are shown in the colored boxes. DNA bulges are indicated as a small additional nucleotide between the normal nucleotides (OT5, between 3 and 4). Numbering on the protospacer starts with the most distal nucleotide on the 5’ end from the PAM sequence as 1. The target adenine is depicted as number 13 within the protospacer. **b,** HTS results to measure editing efficiency for all 35 targetable adenines within the protospacers of five predicted candidate OT loci. HTS was performed for R947X AAV9-control mice and R947X eNme2-C-ABE8e edited mice delivered by AAV9 or AAVMYO (*n*=3 per group). Results for OT editing efficiency are presented for heart, liver, diaphragm, kidney, gastrocnemius (skeletal muscle), spleen and lung. PAM, protospacer adjacent motif; N, (A, T, G, or C). Bars and error bars indicate mean ± s.e.m. Dots represent individual mice.

**Supplementary Fig. 9:**
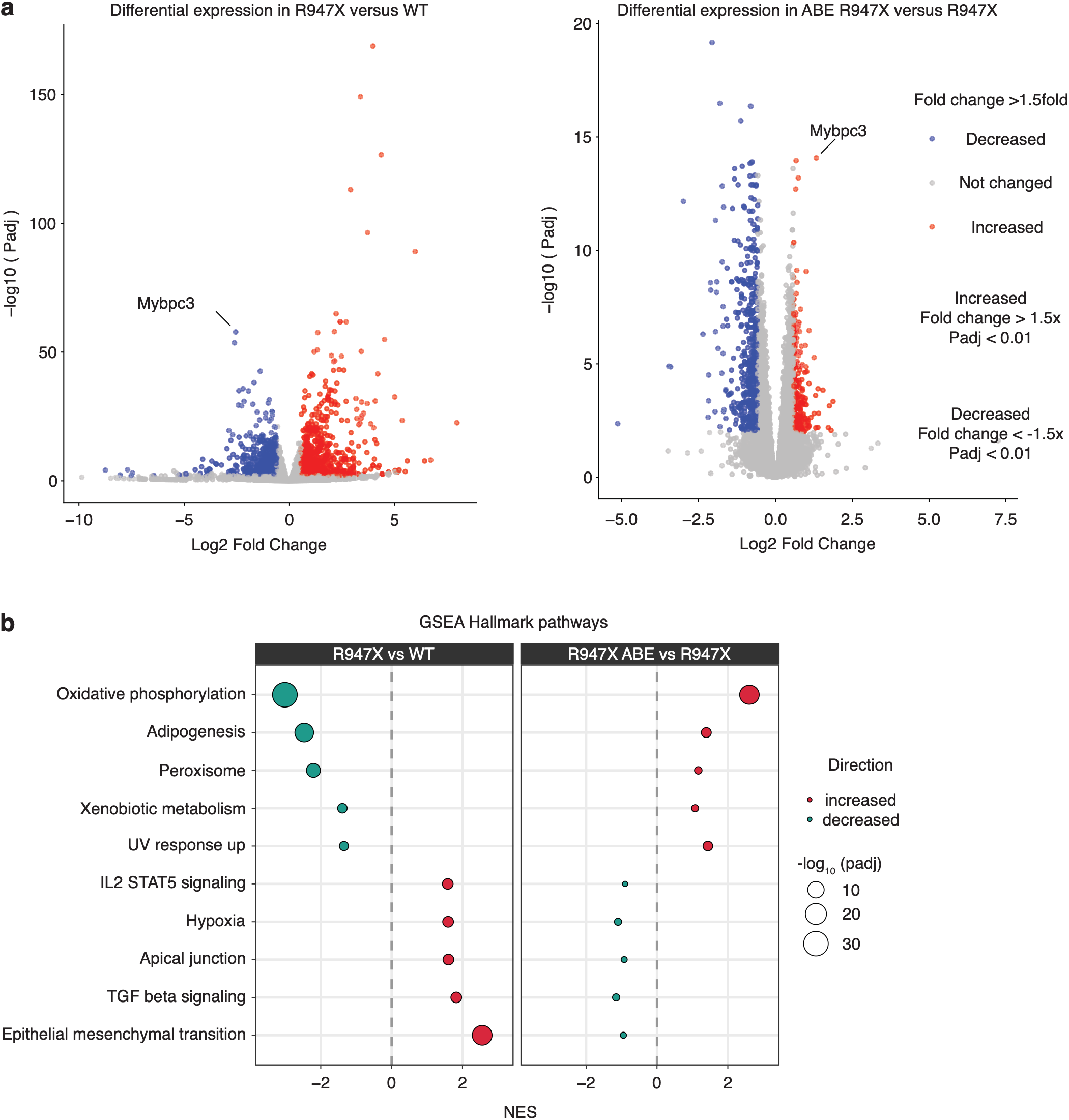
RNA-sequencing analysis of single-AAV ABE in vivo. **a**, Volcano plots showing fold change and P-values of genes up-regulated (red) and down-regulated (blue) in wildtype vs. R947X mice (left) and R947X vs. R947X-ABE edited mice (right). P-adjusted <0.01 and fold change >1.5 were used as cutoffs. P-adjusted values were calculated using Benjamini-Hochberg test. **b**, Gene set enrichment analysis (GSEA) revealing pathways enriched and suppressed in R947X-control vs. wildtype-control and in R947X-edited vs. R947X-control mice at 8 weeks of age (*n*=4 males per group).

## Methods

### hiPSC culturing

hiPSCs were maintained in Essential 8™ Medium (Gibco, A1517001) on Geltrex™ LDEV-Free, hESC-Qualified, Reduced Growth Factor Basement Membrane Matrix (Gibco, A1413302), with medium changes performed daily. For passaging, cells were dissociated with TrypLE Express Enzyme (Gibco, 12605010) and replated in Essential 8™ Medium containing 2 *µ*M Thiazovivin.

### Cardiomyocyte differentiation, expansion and culturing

For cardiomyocyte differentiation, hiPSCs were cultured to 80-90% confluency. On day 0, differentiation was initiated with RPMI-GlutaMAX-HEPES (Gibco, 72400021) supplemented with 4 *µ*M CHIR99021, 0.5 mg/ml human recombinant albumin, and 0.2 mg/ml L-ascorbic acid 2-phosphate. After 48 h, medium was replaced with RPMI-GlutaMAX-HEPES containing 5 *µ*M IWP2, human recombinant albumin, and L-ascorbic acid 2-phosphate for another 48 h. On days 4 and 6, cultures were refreshed with RPMI-GlutaMAX-HEPES containing only human recombinant albumin and L-ascorbic acid 2-phosphate. On day 8, cells were switched to RPMI-GlutaMAX-HEPES supplemented with B-27™ (50x, serum-free; Gibco, 17504001). Cardiomyocyte selection was performed from day 10 till 14 in glucose- and glutamine-free RPMI-1640 (Biological Industries, 01-101-1A) containing B-27™ and 4 *µ*M lactate/HEPES, after which cultures were maintained in RPMI-GlutaMAX-HEPES with B-27™. On day 16, cells were dissociated with TrypLE™ Select Enzyme (Gibco, A1217703) and replated at a 1:10 ratio in RPMI-GlutaMAX-HEPES with B-27™, 2 *µ*M Thiazovivin, and 10% KnockOut™ Serum Replacement (Gibco, 10828028). The following day, cells were expanded for 4 days in RPMI-GlutaMAX-HEPES with B-27™ and 5 *µ*M CHIR99021, then maintained in RPMI-GlutaMAX-HEPES with B-27™ until use.

### Quantitative PCR

Total RNA from hiPSC-CMs and mouse hearts was extracted using TRIzol™ (Thermo Fisher Scientific, 15596018). cDNA was synthesized with the iScript™ cDNA Synthesis Kit (Bio-Rad, 1708891), and qPCR was performed on a Bio-Rad CFX96 Real-Time PCR Detection System using iQ SYBR Green Supermix (Bio-Rad, 1708885). Data were analyzed by the ΔΔCt method, and primer sequences are listed in **Table S1**.

### Immunoblotting

Protein was isolated from hiPSC-CMs and mouse hearts using RIPA lysis buffer (10 mM Tris-HCl, pH 7.5; 150 mM NaCl; 0.1% SDS; 0.5% sodium deoxycholate; 1% Triton X-100) supplemented with cOmplete™ EDTA-free Protease Inhibitor Cocktail (Roche, 11836170001) and PhosSTOP (Roche, 4906837001). Lysates were separated by SDS-PAGE (10-15 *µ*g per lane) and transferred to membranes, which were blocked in 5% milk and incubated with primary antibodies overnight at 4 °C (**Table S8**). HRP-conjugated secondary antibodies (1:10,000; 45 min) were used for detection with the Clarity™ Western ECL Substrate kit (Bio-Rad, 1705061) and imaging on an Amersham Imager 680RGB (GE Healthcare, 29270772). Protein band intensities were quantified using ImageJ.

### Immunofluorescence

For EHMs and mouse heart sections, antigen retrieval was performed by boiling samples for 20 min in Tris-EDTA buffer (10 mM Tris, 1 mM EDTA and 0.05% Tween20 at pH 9.0), followed by cooling for 30 min and washing with PBS. Sections were then incubated for 1 h in blocking buffer (0.5% Triton X-100 and 0.5% BSA in PBS) before overnight incubation at 4°C with the primary antibody diluted in blocking buffer (Table **S8**). After three washes with PBS-T (0.2% Tween20 in PBS), sections were incubated for 1 h with the corresponding secondary antibody (1:200 in blocking buffer) and DAPI. Following three additional washes with PBS-T and one with water, samples were mounted in Mowiol mounting medium (24% w/v glycerol (Baker), 9.6% w/v Mowiol 4-88 (Calbiochem), 0.1 M Tris-HCl, pH 8.5) and imaged using either an Olympus VS200 Research Slide Scanner or a Leica TCS SPE confocal microscope. WGA (Sigma Aldrich, #L4895) staining was used for the analysis of cardiomyocyte cross-sectional area (CSA). CSA was quantified using semi-automated cell segmentations using Cellpose 2.0 and the LabelsToRois plugin in ImageJ.

hiPSC-CMs were cultured on Geltrex-coated glass coverslips or chambered coverslips (ibidi, 81816) and fixed in 4% paraformaldehyde for 30 min at room temperature (RT). After three PBS washes, cells were permeabilized with 0.1% Triton X-100 in PBS for 8 min at RT, blocked for 1 h in 4% goat serum in PBS, and incubated overnight at 4 °C with primary antibodies diluted in the same blocking solution. The coverslips were then washed three times with PBS-T (0.05% Tween20 in PBS) and incubated for 1 h at room temperature (RT) with secondary antibodies (1:200) and DAPI (1:1000). Following three additional PBS-T washes and one water rinse, glass coverslips were mounted in Mowiol and the chambered coverslips were imaged in water. Images were taken using a Leica TCS SPE confocal microscope. Antibodies and the dilutions used can be found in **Table S8**.

### EHM generation

EHM tissues were generated as previously described^62^. Briefly, hiPSC-CMs and human foreskin fibroblasts (HFF) were combined at a 70:30 ratio and resuspended in type I collagen (Collagen Solutions, FS22024) diluted in RPMI 2x (Thermo Fisher Scientific, 51800-035). The cell-collagen suspension was cast into wells of a 48-well EHM plate (myriamed GmbH, myrPlate-TM5) and incubated for 45 min at 37 °C, after which EHM medium freshly supplemented with TGFβ1 was added. For the first 3 days post-casting, tissues were fed daily with TGFβ1-supplemented EHM medium, followed by daily changes with EHM medium alone until day 14. From week 3 onward, cultures were maintained in MEM alpha medium until collection for molecular analyses.

### Contraction analyses

Contraction measurements were obtained by video-optic recording of EHM-induced pole deflection in the myrPlate-TM5 culture format at 37 °C. Spontaneously contracting tissues were recorded for at least 2 min at 50 frames per second at designated time points using a myrImager prototype (myriamed GmbH). The percentage of pole bending was quantified and used as a surrogate measure of contractile force.

### Flow Cytometry

hiPSC-CMs were dissociated using TrypLE™ Select Enzyme (Gibco, A1217703). Cells were centrifuged (300 *g*, 5 min), washed with dPBS, and fixed in ice-cold 70% ethanol. After fixation, cells were permeabilized in blocking buffer (PBS containing 5% fetal bovine serum, 1% BSA, and 0.5% Triton X-100) for 10 min on ice. Cells were incubated with primary antibodies for 1 h at 4 °C, washed with blocking buffer, and subsequently incubated with secondary antibodies for 30 min at RT. After final washes, cells were resuspended in dPBS containing DAPI (1:1000) and analyzed by flow cytometry (BD Biosciences, BD LSRFortessa™ X-20 Cell Analyzer). Antibody details are provided in **Table S8**.

### Mouse fibroblast isolation and immortalization

Liver tissue was finely chopped into small pieces using scissors and transferred into the 5 ml of digestion buffer (2.3ml DMEM/Glu, 2.5 ml dispase, 200 μl liberase TL, 100 μl DNaseI (stock 1 mg/ml)). Samples were incubated at 37 °C in a shaking water bath (180 rpm) for 2 h. Following digestion, the tissue suspension was triturated by pipetting up and down 10 times and passed through a 40 *µ*m cell strainer. Enzyme activity was inhibited by adding 15 ml of PBS supplemented with 10% FBS and centrifuged at 300 *g* for 5 min. The supernatant was aspirated, and the pellet was washed once with 10 ml of PBS supplemented with 10% FBS, followed by centrifugation at 300 *g* for 5 min. Cells were resuspended and plated in DMEM/glucose supplemented with 10% FBS and Primocin (1:500).

Primary fibroblasts were cultured for two weeks prior to immortalization. Cells were then transduced for 24 h with an EF1A-SV40T/CMV-GFP-T2A-puromycin lentiviral vector (VectorBuilder, VB900088-2955hqy). The following morning, the medium was replaced, and transduced fibroblasts were selected with puromycin (2.0 *µ*g/ml, Thermo Fisher Scientific, A1113803) for seven days. Puromycin-containing medium was refreshed on days 2, 4, and 6 and removed on day 7. Following selection, cells were expanded and cryopreserved.

### Mouse embryonic stem cell culture

129/Ola-derived IB10 ES cells (Netherlands Cancer Institute, Amsterdam) were cultured on irradiated mouse embryonic fibroblasts (MEFs) in CM+/+ medium made with GMEM culture media (Thermo Fisher Scientific, 21710-025) supplemented with 1x Glutamax (Thermo Fisher Scientific, 35050-038), 1 mM Sodium Pyruvate (Thermo Fisher Scientific, 11360039), 1x MEM Non-Essential Amino Acids (Gibco, 11140035), 10% FBS, 0.1 mM β-mercaptoethanol and 103u/ml LIF (Millipore, ESG 1107). Cells were dissociated with 10x TVP (dPBS, supplemented with 1 mM EDTA, 1% chicken serum and 0.25% trypsin (Thermo Fisher Scientific, 15090046) and passaged (1:5) onto fresh irradiated MEFs.

### Mouse line generation

sgRNAs were designed using the CCTop - CRISPR/Cas9 target online predictor tool^101^. The selected sgRNA (**Table S9**) was cloned into the pSpCas9(BB)-2A-GFP (PX458) vector backbone (Addgene, #48138). The reference plasmid was generated by cloning a gBlock template (Integrated DNA Technologies; see **Table S9**) containing the mutation of interest into the pUC19 vector backbone (Addgene, #50005). Each gBlock template was flanked by approximately 400 bp homology arms to enable homology-directed repair and included a blocking mutation in the protospacer adjacent motif. This blocking mutation also introduced a restriction enzyme site to facilitate efficient genotyping (**Table S9**). Mouse embryonic stem (mES) cells cultured in a single well of a 12-well plate were dissociated and centrifuged at 300 *g* for 5 min. The supernatant was aspirated, and the cell pellet was resuspended in 500 *µ*l dPBS. Subsequently, 5 *µ*g of the sgRNA plasmid and 5 *µ*g of the reference plasmid were added to the cell suspension. Cells were electroporated using a Bio-Rad Gene Pulser (0.8 kV, 3 *µ*F), incubated at RT for 5 min, and then seeded onto fresh irradiated MEFs. After 24 h, cells were dissociated and single-cell sorted based on GFP expression into MEF-coated 96-well plates using a FACSJazz™ cell sorter (BD Biosciences). After one week, individual clones were dissociated and transferred to two MEF-coated 96-well plates. One plate was used for DNA genotyping using an ethanol-based precipitation method. PCR amplification was performed across the region of interest, followed by restriction enzyme digestion with PvuI-HF (New England Biolabs, R3150) and analysis on a 2% (w/v) agarose gel. Positive clones were further expanded and cryopreserved in liquid nitrogen.

### Karyotyping

To assess genomic DNA integrity of successfully modified mES cells, cultures were treated with 0.05 *µ*g/ml colcemid (Gibco, 152120012) for 90 min at 37 °C. Medium was collected, cells were rinsed once with dPBS, dissociated with 10× TVP for 5 min, and centrifuged at 300 *g* for 5 min. The cell pellet was resuspended in 1 ml of 75 mM KCl prewarmed to 37 °C and incubated for 10 min at 37 °C with gentle shaking. Fixation was initiated by adding 50 *µ*l of freshly prepared methanol:acetic acid (3:1) dropwise, followed by centrifugation at 300 *g* for 5 min. The supernatant was removed, and cells were fixed in 1 ml methanol:acetic acid (3:1) added dropwise while gently shaking, followed by a 20-minute incubation at 37 °C. Cells were centrifuged at 300 *g* for 5 min, and the fixation step was repeated twice. After the final centrifugation, the pellet was resuspended in 0.5 ml methanol:acetic acid (3:1). Cell suspensions were dropped from approximately 50 cm onto glass microscope slides and air-dried. Slides were mounted with ProLong™ Gold Antifade Mountant with DAPI (Thermo Fisher Scientific, P36935). Chromosome spreads were imaged using a Leica TCS SPE confocal microscope. Clones with ≥80% of metaphase spreads displaying a normal karyotype (40 chromosomes) were selected for blastocyst injection.

### Echocardiographic analysis

Cardiac function and dimensions were assessed by two-dimensional transthoracic echocardiography using a Vevo® 2100 ultrasound system (VisualSonics) equipped with a 30 MHz transducer. Mice were sedated with 2% isoflurane and placed on a heated platform. The heart was imaged in parasternal long-axis and short-axis views at the level of the papillary muscles. M-mode recordings obtained from the long-axis view were used to measure left ventricular mass (LV mass), left ventricular end-diastolic diameter (diameter; d), left ventricular end-systolic diameter (diameter; s), left ventricular anterior wall thickness at end diastole (LVAW; d) and end systole (LVAW; s), left ventricular end-diastolic volume (volume; d), left ventricular end-systolic volume (volume; s), ejection fraction (EF) and fractional shortening (FS). For each parameter, at least six measurements were averaged per animal. Fractional shortening (FS) and ejection fraction (EF), automatically calculated using Vevo® LAB software version 1.7.1, were used to assess cardiac contractile function.

### Tissue Collection

Following echocardiographic assessment, mice remained under isoflurane anesthesia and were euthanized by cervical dislocation. Tissues were collected, rinsed in ice-cold PBS, weighed, and either snap-frozen in liquid nitrogen for protein and RNA analyses or fixed in 4% paraformaldehyde for histological analysis. The left hind leg was enzymatically digested with proteinase K (final concentration 0.6 *µ*g/*µ*l) in lysis buffer, after which tibia length (TL) was measured using a digital caliper.

### Lentivirus production and transduction

HEK293T cells were seeded to reach 80-100% confluency on the day of transfection. Cells were co-transfected using polyethyleneimine (PEI) with the transfer plasmid, pMD2.G (Addgene, #12259), and psPAX2 (Addgene, #12260). After 6-8 hours, the medium was refreshed. 48 hours after transfection, viral supernatant was collected, centrifuged at 300 *g* for 5 min, and filtered through a 0.45 *μ*m membrane. Lentiviral particles were aliquoted and stored at -80 °C until use. For high-titer preparations, lentivirus was concentrated using the Lenti-X™ Concentrator (Takara, #631231) according to the manufacturer’s instructions. Briefly, viral supernatant was collected, centrifuged, and transferred to a sterile tube. Concentrator was added at a 1:3 ratio and mixed until homogeneous, followed by incubation at 4 °C for at least 30 min and up to 7 days. Samples were then centrifuged at 1,500 *g* for 60 min, the supernatant was discarded, and viral pellets were resuspended in dPBS, aliquoted, and stored until use.

### HEK293T-R943X cell line generation

HEK293T cells (80-100% confluent) were transduced in 6-well plate with lentivirus containing the target R943X sequence followed by a hygromycin cassette. The following morning, the medium was replaced, and transduced cells were selected with Hygromycin B (200 *µ*g/ml, Thermo Fisher Scientific, 10687010) for seven days.

Hygromycin B containing medium was refreshed on days 2, 4, and 6 and removed on day 7. Following selection, cells were expanded and cryopreserved.

### AAV injections in mice

AAV vectors encoding the different promoters driving eGFP expression and the adenine base editors were generated in collaboration with the Giacca and Zentilin laboratories (Trieste, Italy). Pups were IP injected at P3 using a 30-gauge needle fitted to an insulin syringe.

### HTS of on- and off-target PCR-amplified target sites

gDNA from in vitro experiments was isolated using the NucleoSpin Tissue kit (MACHEREY-NAGEL, 740952), DirectPCR Lysis Reagent (Gentaur, 0388-301-C) and gDNA from mouse tissue were isolated using the DNeasy Blood & Tissue Kit (Qiagen, 69506) according to the manufacturer’s protocol. PCR1 reactions used Q5® Hot Start High-Fidelity 2× Master Mix (New England Biolabs, M0494) with 50 ng input gDNA or 4 *µ*l lysate (when using DirectPCR lysis Reagent) in 25 *µ*l reactions according to the manufacturer’s protocol. Primers contained custom barcodes and partial Illumina adapters. PCR1 was performed under the following cycle conditions: 98 °C for 30 s, (98 °C for 10s, 65 °C for 30s, 72 °C for 30s) x 34 cycles, 72 °C for 5 min. After PCR1, samples were purified by electrophoresis on a 2% agarose gel using the NucleoSpin Gel and PCR Clean-up kit (MACHEREY-NAGEL, 740609) and sequenced at Azenta Genewiz (Illumina® MiSeq platform). Samples sent to USEQ were prepared using a PCR2 reaction. In short, 2 *µ*l of PCR1 reaction volumes were pooled and purified using CleanNGS magnetic beads (CNGS-0050) (0.8x beads to PCR volume ratio). PCR2 was performed to add Illumina flow cell binding sequences and indexes. PCR2 used Q5® Hot Start High-Fidelity 2× Master Mix (New England Biolabs, M0494) and all PCR2 reactions were performed using the following cycling conditions: 98 °C for 30 s, (98 °C for 10s, 65 °C for 30s, 72 °C for 30s) x 8 cycles, 72 °C for 5 min (100 ng input DNA). PCR2 libraries were purified by electrophoresis on a 2% agarose gel using the NucleoSpin Gel and PCR Clean-up kit (MACHEREY-NAGEL, 740609). PCR2 library concentrations were quantified using the Qubit™ dsDNA High Sensitivity Assay Kit (Thermo Fisher Scientific, Q32851). PCR2 amplicon libraries were diluted to 4 nM, pooled and sequenced at the Utrecht Sequencing Facility USEQ (Illumina® NovaSeq X, 300 cycles, 2×150 bp). All primers are listed in **Table S6**. sgRNA-dependent DNA off-targets were computationally selected using Cas-OFFinder^100^.

### HTS data analysis

Samples were demultiplexed using the Ultraplex package from the Ulelab with parameters -m5 0 and -q 30. Sequencing data were analyzed using CRISPResso2 (version: 2.1.3) in BE batch mode with the following parameters: -w12, -wc -12, -q 30^102,103^. Base editing percentages were calculated from the specified base conversion in the genomic sequence, with background editing corrected against the corresponding control samples, using the Nucleotide_percentage_summary.txt output file.

### Viral copy number assay

Genomic DNA was isolated using the DNeasy Blood & Tissue Kit (Qiagen, 69504). AAV vector genomes were determined by PrimeTime^TM^ qPCR Probe assays using the PrimeTime^TM^ Gene Expression Master Mix (IDT, 1055770). Custom primers and probes were designed using the PrimerQuest Tool (IDT) (**Table S4**). The primers and probes anneal to the 6C2 promoter (ABE studies) and to a U6 sgRNA cassette (promoter studies). The AAV viral copy number was normalized to the mouse glucagon gene copy number as previously described^99^.

### Promoter development and validation

Chimeric promoter sequences were generated using a custom-built python script. Novel promoters were synthesized as pooled oligos and subsequently cloned into an AAV-backbone plasmid using Gibson assembly (NEBuilder HiFi DNA Assembly master Mix; NEB, E2621L). Following transformation, 1,000 unique promoter plasmids were isolated and verified using Sanger sequencing. For validation in vitro, hiPSC-CMs were transfected as previously described^10^. In short, hiPSC-CMs were plated in 96-well glass bottom microplates (Corning, 4580) coated with Matrigel (Corning, 356231) and transfected ten days after seeding using 100 ng of plasmid DNA in OPTI-MEM^TM^ (Thermo Fisher) per well. ViaFect (Promega, E4983) was used according to the manufacturer’s protocols. Promoter activity was assessed by measuring GPF intensity six days after transfection using Cytation 5 imaging. Transfected hiPSC-CMs were considered positive or GPF expression above the arbitrary intensity minimum of 250 (a.u). The 400 bp promoter cTnT promoter library served as positive control.

For in vivo validation, mice and NHPs were IV-injected at 8 weeks and 5-6 years of age, respectively. DNA and RNA from heart and liver was isolated using the DNeasy Blood & Tissue Kit (Qiagen, 69506) and the RNeasy Mini Kit (Qiagen, 74106). Next, barcoded regions of DNA and cDNA (Invitrogen SuperScript III First-Strand Synthesis SuperMix) were PCR-amplified and sent for HTS. For data analysis, raw barcode counts of DNA and cDNA were normalized to total sequencing depth, the abundance of the barcode in the initial virus pool and the measurement of the reference cassette within the pool. Counts for all three barcodes associated with each promoter construct were averaged. As last, the relative cDNA counts were normalized to the viral DNA counts in the same sample to generate the activity score for each promoter.

### Molecular cloning

The pCMV-eNme2-C-ABE8e plasmid (Addgene, #185566) and pAAV-EFS-SAKKH-ABE8e-bGH-U6-sgRNA-BsmBI (Addgene, #189923) were used to construct the base editors and clone them into a lentiviral backbone (Clontech, #632185) by Gibson assembly of PCR-amplified fragments or by standard restriction enzyme cloning. ABE9 and MYBPC3 variants were generated by Gibson assembly using gene fragments synthesized by Integrated DNA Technologies. sgRNAs (**Table S7**) were cloned with Gibson assembly using bridge oligos or by Golden Gate assembly. For promoter studies, either eGFP or Cas9 (Addgene, #48138) expression was driven by the different promoters; promoter sequences are provided in **Table S3**. All plasmids were purified using the PureLink™ Quick Plasmid Miniprep Kit (Thermo Fisher Scientific, K210011). For high-yield plasmid preparations, midi- and maxi-preps were performed using the PureLink™ HiPure Plasmid Midiprep Kit (Thermo Fisher Scientific, K210004) and PureLink™ HiPure Plasmid Maxiprep Kit (Thermo Fisher Scientific, K210007), respectively.

### RNA-sequencing and data analysis

Total RNA was extracted from wildtype control, R947X control and AAVMYO-treated mice (n=4 per group) using standard TRIzol extraction (Thermo Fisher). Illumina TruSeq stranded mRNA libraries were sequenced paired 2×75 base pairs on Illumina NovaSeq X. Reads were mapped to mouse genome (GRCm38) and differential expression analysis was performed using DESeq2 1.50.2. Differentially expressed genes were those with absolute fold change of 1.5 and adjusted Benjamini-Hochberg adjusted P < 0.01. Functional annotation was performed using fGSEA 1.36.2 with molecular signatures database (MsigDB mouse version 2026.1)^104^.

### MYBPC3 protein structures

Protein structure figures were generated using PyMOL (Schrödinger, LLC; https://www.pymol.org/).

### Quantification and statistical analysis

Data are reported as mean ± s.e.m. and analyzed with GraphPad Prism 11. Outliers were removed via the ROUT method (Q=1%). Data normality was evaluated using the Kolmogorov-Smirnov test. Comparisons between two groups were analyzed using a two-tailed *t*-test (two groups) or one-way ANOVA (multiple groups). Non-parametric data were analyzed using Mann-Whitney or Kruskal-Wallis tests. Multiple comparison to a control condition for one-way ANOVA was performed with Dunnett’s test.

### Ethical approval

All animal studies carried out at the Hubrecht Institute adhered to the institutional guidelines and complied with the regulations set forth by the Animal Welfare Committee of the Royal Netherlands Academy of Arts and Sciences. Animal experiments were conducted upon approval by the ‘Animal Welfare Body Utrecht’ (I.v.D.) of the Royal Dutch Academy of Sciences and Arts (K.N.A.W.) and are in compliance with national legislation and institutional guidelines. Male and female mice were included in the studies performed. Mouse lines were maintained on a C57B/6J background.

All animal procedures performed in the USA were reviewed and approved before study initiation by the Institutional Animal Care and Use Committee (IACUC) at the respective institutions. Mouse studies were conducted at Tenaya Therapeutics in South San Francisco, CA, USA, under approved protocol, 2023.007. Non-human primate studies were conducted at Altasciences Preclinical Columbia in Auxvasse, MO, USA, under an approved protocol. Animal care and experimental procedures complied with the “Guide for the Care and Use of Laboratory Animals, 8th edition,” published by the US National Institutes of Health (ISBN10: 0-309-15400-6).

## Acknowledgements

We thank Greenstone Biosciences for providing the patient-derived hiPSCs used in this study. We thank the Utrecht Sequencing Facility (USEQ) for providing sequencing service and data. USEQ is subsidized by the University Medical Center Utrecht and The Netherlands X-omics Initiative (NWO project 184.034.019). We thank Bram Verhagen for his invaluable advice on the design of HTS sequencing libraries. We thank Jeroen Korving for his help with preparing the slides for mouse tissue imaging experiments. We thank everyone from our Flow Cytometry Facility at the Hubrecht Institute for helping with sorting experiments.

## Author contributions

Conceptualization: T.M., S.H., M.A.M., C.A.R., C.J.B. and E.v.R. Investigation: T.M., S.H., L.S., S.K., C.A., H.d.R., A.L., T.W., M.C., I.P., D.V., M.A.M., C.A.R., P.R.K. Resources: A.E.d.L., M.W.C., S.J.v.K., M.W., L.Z., D.D.S., K.N.I., M.G. Data analysis: T.M., S.H., L.S., S.K., P.R.K., M.A.M., C.J.B. Funding acquisition: E.v.R. Writing - original draft: T.M., C.J.B., E.v.R. Writing - review and editing: all authors.

## Funding

This work was supported by the Foundation Leducq Transatlantic Network of Excellence (17CVD02 to M.G. and E.v.R.) and the REPAIR project (2023TKI2410), funded through the Public-Private Partnership (PPP) Allowance made available by Health Holland, Top Sector Life Sciences & Health, through University Medical Center Utrecht, with co-funding from Tenaya Therapeutics.

## Ethics declaration

E.v.R. is a consultant for Tenaya Therapeutics and Novo Nordisk and is Chief Scientific Officer of Phlox Therapeutics and Chief Executive Officer of Revier Therapeutics. M.G. is founder, consultant, member of the Board and equity holder in Purespring Therapeutics, Forcefield Therapeutics and Heqet Therapeutics. K.N.I. is an employee and a shareholder of Tenaya Therapeutics. P.R.K. is a shareholder of Tenaya Therapeutics. The other authors declare no competing interests.

## Notes

### Competing Interest Statement

E.v.R. is a consultant for Tenaya Therapeutics and Novo Nordisk and is Chief Scientific Officer of Phlox Therapeutics. M.G. is founder, consultant, member of the Board and equity holder in Purespring Therapeutics, Forcefield Therapeutics and Heqet Therapeutics. K.N.I. is employee and shareholder of Tenaya Therapeutics. P. R. K. is shareholder of Tenaya Therapeutics. The other authors declare no competing interests.

